# Normalization benchmark of ATAC-seq datasets shows the importance of accounting for GC-content effects

**DOI:** 10.1101/2021.01.26.428252

**Authors:** Koen Van den Berge, Hsin-Jung Chou, Hector Roux de Bézieux, Kelly Street, Davide Risso, John Ngai, Sandrine Dudoit

**Affiliations:** Department of Statistics, University of California, Berkeley, CA, USA; Department of Applied Mathematics, Computer Science and Statistics, Ghent University, Ghent, Belgium; Bioinformatics Institute Ghent, Ghent University, Ghent, Belgium; Department of Molecular and Cell Biology, University of California, Berkeley, CA, USA; Division of Biostatistics, School of Public Health, University of California, Berkeley, CA, USA; Center for Computational Biology, University of California, Berkeley, CA, USA; Department of Data Sciences, Dana-Farber Cancer Institute, Boston, MA, USA; Department of Biostatistics, Harvard T.H. Chan School of Public Health, Boston, MA, USA; Department of Statistical Sciences, University of Padova, Italy; Helen Wills Neuroscience Institute, University of California, Berkeley, CA, USA

## Abstract

Modern assays have enabled high-throughput studies of epigenetic regulation of gene expression using DNA sequencing. In particular, the assay for transposase-accessible chromatin using sequencing (ATAC-seq) allows the study of chromatin configuration for an entire genome. Despite the gain in popularity of the assay, there have been limited studies investigating the analytical challenges related to ATAC-seq data, and most studies leverage tools developed for bulk transcriptome sequencing (RNA-seq). Here, we show that GC-content effects are omnipresent in ATAC-seq datasets. Since the GC-content effects are sample-specific, they can bias downstream analyses such as clustering and differential accessibility analysis. We introduce a normalization method based on smooth-quantile normalization within GC-content bins, and evaluate it together with eleven different normalization procedures on eight public ATAC-seq datasets. Our work clearly shows that accounting for GC-content effects in the normalization is crucial for common downstream ATAC-seq data analyses, leading to improved accuracy and interpretability of the results. Using two case studies, we show that exploratory data analysis is essential to guide the choice of an appropriate normalization method for a given dataset.

## 1 Introduction

Genomic DNA is packaged into chromatin in the eukaryotic nucleus via a highly-deliberate and dynamic process. The study of chromatin configuration aids in unraveling the complex epigenetic regulation of gene expression. Chromatin accessibility, which reflects the relatively open or closed chromatin conformation, affects the ability of nuclear proteins to physically interact with chromatin DNA and hence regulate gene expression ^1^. Genome-wide mapping of chromatin accessibility delineates the functional chromatin landscape corresponding to transcription start sites (TSS), transcription factor (TF) binding sites, and all classes of cis-regulatory elements (e.g., promoters and enhancers) ^2,3^. ATAC-seq, a robust assay for transposase-accessible chromatin using sequencing, has been used to provide insight into chromatin accessibility with a relatively simple and time-saving protocol ^1,4^.

ATAC-seq relies on a hyperactive Tn5 transposase that can simultaneously cut accessible DNA fragments and ligate sequencing adapters to both strands ^4^. The tagged DNA fragments are amplified, sequenced, and mapped back to the genome, upon which accessible regions are identified by the enrichment of mapped read ends, traditionally using peak-calling algorithms ^5,6^. The data used for downstream analysis thus typically consist of a count matrix, where each row corresponds to a genomic region (or “peak”) and each column corresponds to a sample. The count in each cell of the matrix represents the number of read ends mapped to a particular peak for a given sample, and is a proxy for the accessibility of the genomic region.

High-throughput sequencing studies are often influenced by several factors of technical variation, e.g., sample preparation, library preparation, and sequencing batch ^7,8^. Notably, GC-content, the fraction of guanine and cytosine nucleotides in a particular genomic region or gene, has previously been identified as a sample-specific technical bias factor, e.g., in peak-calling for chromatin immunoprecipitation sequencing (ChIP-seq) data ^9^ or normalization and differential expression for RNA-seq data ^10,11^. Similarly, ATAC-seq data have been shown to be affected by technical variation due to, for example, enzymatic cleavage effects, PCR bias, and duplicate reads ^12^. Importantly, the effect of GC-content in ATAC-seq data has been noted previously ^13,14^. Indeed, since a large fraction of accessible regions are around gene promoters, which often have a high GC-content and are enriched in CpG islands ^15^, we naturally expect an association between accessibility and GC-content. However, the impact of GC-content effects in ATAC-seq data on downstream analyses has not been studied in depth.

Notwithstanding the recent surge in popularity for the assay, surprisingly few studies investigate the analytical challenges of ATAC-seq data, e.g., normalization and differential accessibility (DA) analysis. Indeed, most data analysis workflows rely on statistical methods originally developed for ChIP-seq or bulk RNA-seq data to analyze bulk ATAC-seq datasets ^16–18^. Recently, Reske et al. ^18^ compared pipelines for DA analysis and showed that normalization has a large influence on the results. While the authors advised comparing multiple normalization methods for a particular dataset at hand, they did not elaborate on GC-content effects nor normalization methods that take this into account. In particular, while most research papers analyzing bulk ATAC-seq data adopt standard bulk

RNA-seq global-scaling normalization procedures (e.g., Rizzardi et al. ^16^, Philip et al. ^17^), such as total-count normalization, edgeR’s trimmed mean of *M* -values (TMM) ^19^, or DESeq2’s median-of-ratios (MOR) ^20^, some account for GC-content effects ^14^, typically through conditional-quantile normalization (cqn) ^21^. Given this dichotomy in normalization choices, we investigate the influence of accounting for possible GC-content effects on downstream analyses for ATAC-seq data.

In this manuscript, we show that GC-content effects in ATAC-seq data can be sample-specific, which indicates that they can bias downstream analyses such as clustering and differential accessibility analysis. We introduce a normalization method, smooth GC-FQ, based on smooth-quantile normalization within ^22^ GC-content bins, and evaluate it together with several GC-aware as well as GC-unaware normalization methods using a principled framework. We further study the impact of GC-content effects on the accuracy and interpretation of differential accessibility analysis results. While no normalization method uniformly performs best across all datasets, smooth GC-FQ performs best on average, and GC-aware normalization methods typically perform better than GC-unaware methods, emphasizing the need to correct for GC-content effects. We recommend that researchers use exploratory data analysis methods to guide the choice of normalization method.

## 2 Results

### 2.1 GC-content effects are sample-specific and confound downstream analyses

In ATAC-seq, accessibility counts are often positively associated with GC-content. We explore this using data from Calderon et al. ^23^, who generated an ATAC-seq atlas of human immune cell types. Here, we focus on six replicate samples of memory natural killer (MNK) cells in a control condition.

Figure 1a shows that the accessibility count of a particular genomic region is associated with its GC-content. However, the slope and shape of the curves may differ between samples, which indicates that GC-content effects are sample-specific and can therefore bias between-sample comparisons. This can be similarly observed for other cell types in this dataset (Supplementary Figure 1). Note that, while the width of each peak is also associated with its accessibility, this effect tends to be more uniform across different samples and therefore have lower impact on between-sample comparisons (Supplementary Figure 2). This could be considered analogous to the effect of gene length on read counts in RNA-seq data.

**Figure 1:**
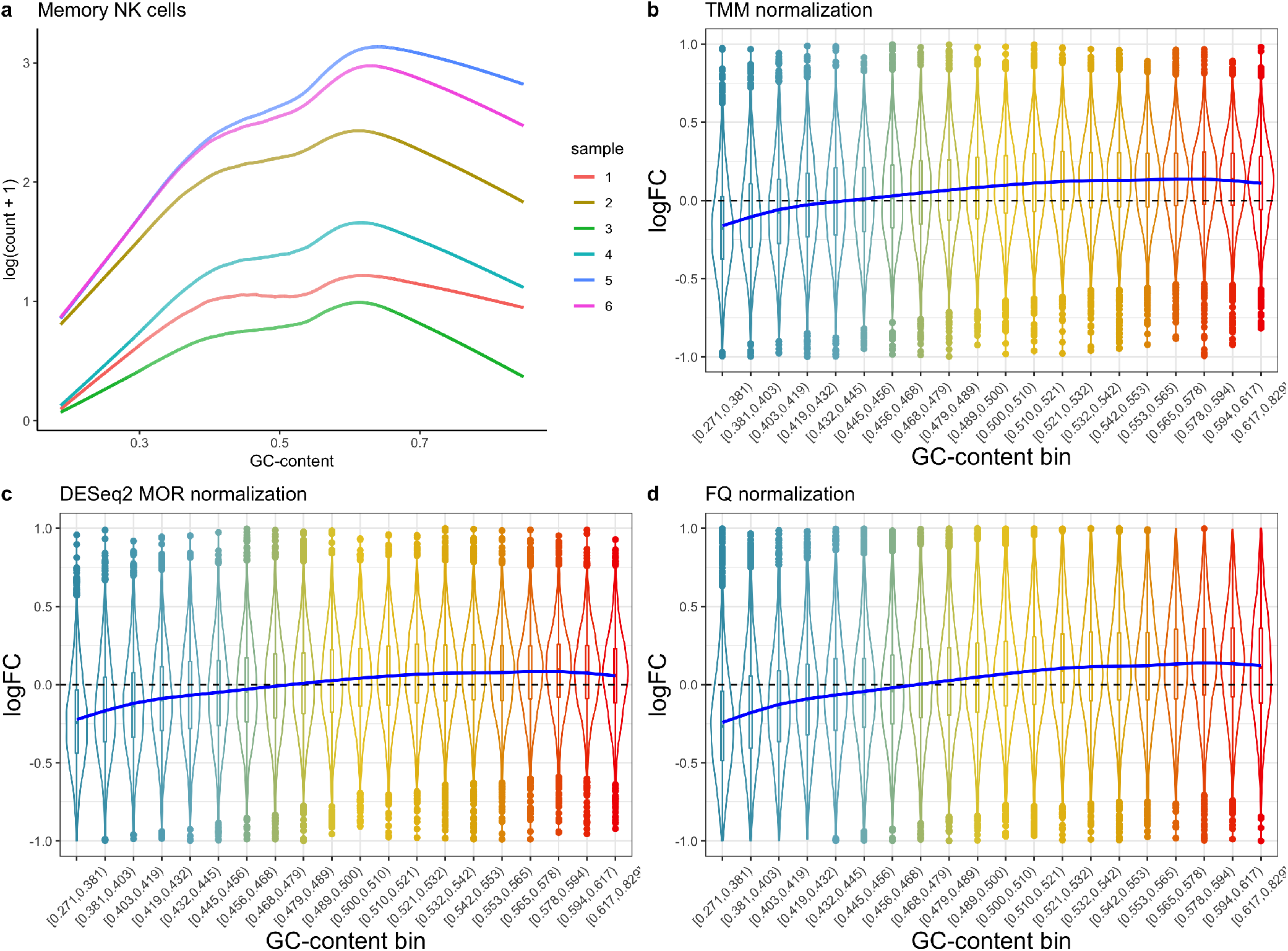
GC-content effects are sample-specific and confound differential accessibility analysis. (a) Fitted lowess curves of log-count as a function of GC-content for the six MNK cell control samples in Calderon et al. ^23^. The shape and slope of the curves can be different for different samples, especially Sample 1 in comparison to other samples. This is also reflected in the data for other cell types (Supplementary Figure 1). (b) Differential accessibility log-fold-changes for a 3 vs. 3 mock null comparison, based on normalization and differential accessibility analysis using edgeR, show a bias for peaks with low and high GC-content (in a null setting, LFC should be centered around zero). The blue curve represents a generalized additive model (GAM) fit. (c) Similar to (b), but using DESeq2 for normalization and differential accessibility analysis. (d) Similar to (b), but using full-quantile normalization and edgeR differential accessibility analysis.

One might initially think that, because DA analyses involve comparing read counts between samples for a given genomic region with a fixed GC-content, GC-content effects would cancel out. However, because of their sample-specificity, GC-content effects also impact log-fold-changes (LFC) comparing accessibility between samples for a given region. A 3 vs. 3 mock null comparison of the MNK cells (i.e., a comparison of the same type of cells which should not exhibit DA), using both edgeR ^24^ and DESeq2^20^, reveals a bias in the LFC with respect to GC-content (Figure 1b-c). That is, the LFC are not centered around zero, as expected for a null comparison of normalized data, and also vary with GC-content. Furthermore, both TMM and DESeq2 normalizations, which are frequently used for DA analysis in ATAC-seq data, fail to remove GC-content effects. Full-quantile (FQ) normalization ^25^, another popular normalization method, also fails at removing GC-content effects (Figure 1d). Similar effects can be observed for other cell types for which six replicates are available in the same condition (Supplementary Figures 3-4).

If not accounted for, GC-content effects can have a significant impact on a downstream differential accessibility analysis, masking biological signal and also leading to false positives, as was similarly previously observed in RNA-seq data ^10,11,21^

### 2.2 GC-aware normalization

GC-content effects have been observed and accounted for in other work on ATAC-seq data ^14,26^, typically using conditional-quantile normalization (cqn) ^21^ (see Methods). While this method removes GC-content effects in many datasets, Risso et al. ^10^ observed that, in the context of RNA-seq, cqn’s regression approach may be “too weak” for some datasets and “more aggressive normalization procedures” may be required. They proposed a method, implemented in the Bioconductor R package EDASeq, based on two rounds of full-quantile normalization: First, FQ normalization within a sample across bins of features (e.g., peaks or genes) with similar GC-content and, subsequently, FQ normalization between samples. We will denote this method as FQ-FQ normalization.

However, read count distributions may in some datasets be more comparable across samples within a GC-content bin than across GC-content bins within a sample (see Figure 2a). This motivates a variant of FQ-FQ normalization, which we call GC-FQ, that applies full-quantile normalization across samples for each GC-content bin separately.

**Figure 2:**
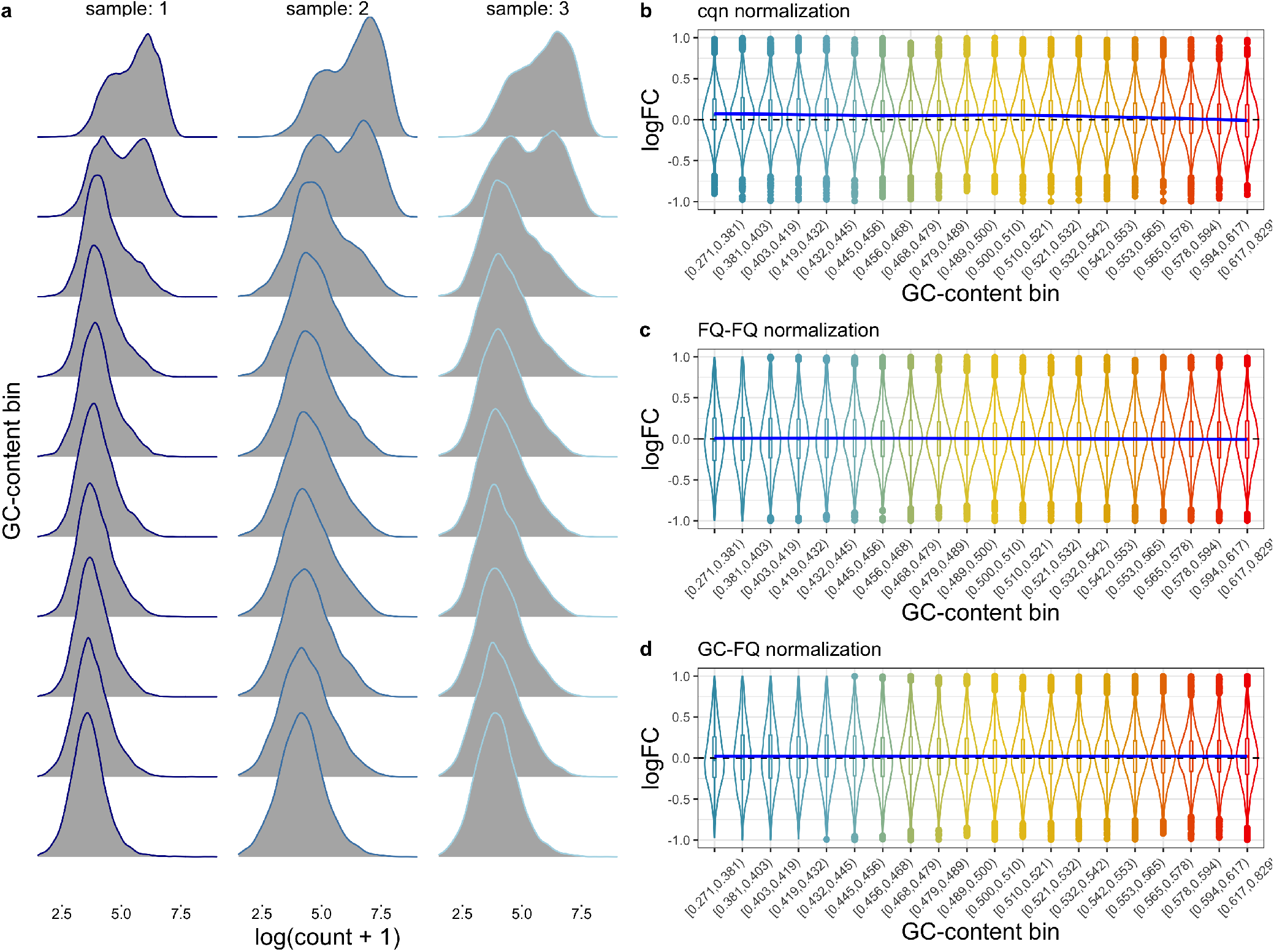
GC-aware normalization methods cqn, FQ-FQ, and GC-FQ are successful in eliminating GC-content effects on the differential accessibility fold-change estimates. (a) Accessibility distributions for three replicates from the dataset of Philip et al. ^17^. The peaks are grouped into 10 equally-sized bins according to their GC-content (rows) and the accessibility distribution (kernel density estimate) is plotted for each bin. The distributional shapes are more comparable across samples for a particular GC-content bin, than they are across GC-content bins for a particular replicate. (b-d) There is no visible GC-content effect on fold-changes estimated using edgeR following normalization with GC-aware methods cqn, EDASeq, and GC-FQ, in the mock comparisons for the dataset from Calderon et al. ^23^. The blue curve represents a GAM fit.

Note that this attempts to equalize the GC-content effect across samples and therefore performs between-sample normalization simultaneously.

For the ATAC-seq dataset of Calderon et al. ^23^, all three GC-aware methods (cqn, GC-FQ, and FQ-FQ) indeed effectively remove GC-content effects (Figure 2b-d) on the fold-changes.

All three of these GC-aware normalization methods rely on full-quantile between-sample normalization, which is an aggressive normalization method that does not come without assumptions. Since FQ normalization forces distributions to be equal across all samples, the underlying assumption is that global differences between distributions are the result of technical effects. In other words, if there were no technical nor sampling variability in the data, the distributions of all samples should be identical, hence comparable, and this is what FQ normalization is trying to achieve. This assumption is restrictive and is not guaranteed to hold for all datasets. Hicks et al. ^27^ recently developed a generalization of FQ normalization, smooth-quantile normalization, or qsmooth, that can account for global differences between distributions due to biological effects of interest. The method is based on the assumption that the read count distribution of each sample should be equal within biological groups or conditions, but could vary between groups. Essentially, the method is a weighted combination of FQ normalization between samples for each biological group separately and FQ normalization across all samples of all biological groups.

We therefore also implement a variant of GC-FQ, which we call smooth GC-FQ, that applies smooth-quantile normalization across samples within each GC-content bin separately and is therefore capable of dealing with biological groups that have global distributional differences between them.

### 2.3 Benchmarking ATAC-seq normalization

We evaluate normalization methods using eight public bulk ATAC-seq datasets ^14,17,23,28–32^, including ATAC-seq atlases of mouse tissues ^30^, human blood cells ^23^, and human brain cells ^29^, thus spanning a multitude of biological systems. For each dataset, we use the publicly-available raw (i.e., unnormalized) accessibility count matrix; the different datasets hence also span a realistic range of preprocessing and peak-calling pipelines. We compare GC-aware normalization methods (smooth) GC-FQ, cqn, and FQ-FQ, with GC-unaware normalization methods qsmooth, TMM, DESeq2, full-quantile, total-count, upper-quartile, and no normalization; see Methods for a description of each normalization procedure.

Our benchmarking framework evaluates different aspects of each normalization method, which can broadly be categorized as follows: (i) between-sample comparison of normalized expression measures, (ii) performance in differential accessibility analysis, (iii) removal of GC-content effects, each of which are described in more detail below. The specific measures used and their definitions for each of these components are described in Supplementary Methods.

### Between-sample comparison of normalized expression measures

We evaluate and rank normalization methods based on a range of performance measures implemented in scone ^33^. While scone provides a valuable framework for benchmarking normalization procedures, we find that some default measures may favor certain normalization methods over others. We therefore use simulated mock datasets as well as real datasets to select relevant measures for our context, as described in Supplementary Results. Based on this evaluation, we benchmark normalization methods using five summary measures. The first three are the average silhouette width for (i) a clustering of samples according to biological covariate(s) of interest (*Bio Sil*); (ii) a clustering of samples according to (unwanted) batch effects (*Batch Sil*); (iii) an empirical clustering of samples using partitioning around medoids (*PAM Sil*). We also evaluate normalized data by the correlation of their log-count principal components with (iv) principal components of QC variables (see Methods section for which QC variables are used in each dataset), and (v) principal components of factors of unwanted variation, derived from negative control features. Here, we use peaks that overlap with housekeeping genes as negative control features.

### Performance in differential accessibility analysis

The differential accessibility analysis performance evaluation relies on two scenarios, based on synthetic null and synthetic signal datasets.

First, a mock null analysis is performed for each real dataset where, for each stratum of the biological covariate of interest, samples are split randomly into two groups to create a mock variable. Since the two mock groups therefore contain a similar number of samples from each stratum, we expect no systematic differences between the groups. A differential accessibility analysis is then performed using each of the normalization procedures (see Methods). The following two evaluation measures are computed: The fraction of peaks returned at a nominal marginal significance level of 5% (*FPR*) and the Hellinger distance of the *p*-value distribution with a uniform distribution on the interval [0, 1] (*P-val unif*). Both measures aim to assess control of false positives in a DA analysis.

Second, we use each real dataset to construct synthetic signal datasets of 12 samples each, based on the simulation framework described in Methods. We use the simulated datasets to assess DA analysis performance based on the area under the receiver operating characteristic (AUROC) curve (*auroc*).

### GC-content effect removal

Finally, we use the evaluations in both components above in combination with three measures that assess the removal of GC-content effects. In the scone normalization performance evaluation, we use a measure based on relative log-expression (RLE) values ^34^ to investigate whether the normalization works across the range of GC-content values (*RLE GC*), see Supplementary Methods for details. In the mock comparison, we assess deviation of *p*-value uniformity as a function of GC-content, by calculating the variability in Hellinger distance between the *p*-value distributions in each of 20 equally-sized GC-content bins and a uniform distribution. Good normalization methods should have a similar *p*-value distribution across GC-content bins (*p-val GC*). Finally, we use the DA analysis on the simulated datasets to calculate the distance in empirical cumulative distribution functions between the observed GC-content distribution of called DA peaks and the GC-content distribution of truly DA peaks (*GC-dist DA*).

The benchmark results for each dataset are shown in Supplementary Figure 5, and summarized across datasets in Figure 3. While no method uniformly outcompetes all others, smooth GC-FQ performs best in 7 out of 8 datasets. Other GC-aware normalization methods, such as FQ-FQ and GC-FQ, also often perform well, while the performance of cqn is variable across datasets. GC-unaware normalization methods typically perform worse than GC-aware methods. Out of the former, qsmooth and FQ consistently perform reasonably well. The good performance of both smooth GC-FQ and qsmooth suggests that, in bulk ATAC-seq data, often large numbers of differentially abundant features between biological conditions may exist, possibly more so than what is typically observed in bulk RNA-seq data. Indeed, this is also what we observe in the two case studies discussed below, where, for both datasets, many methods flag over 35% of all features as differentially accessible between biological conditions.

**Figure 3:**
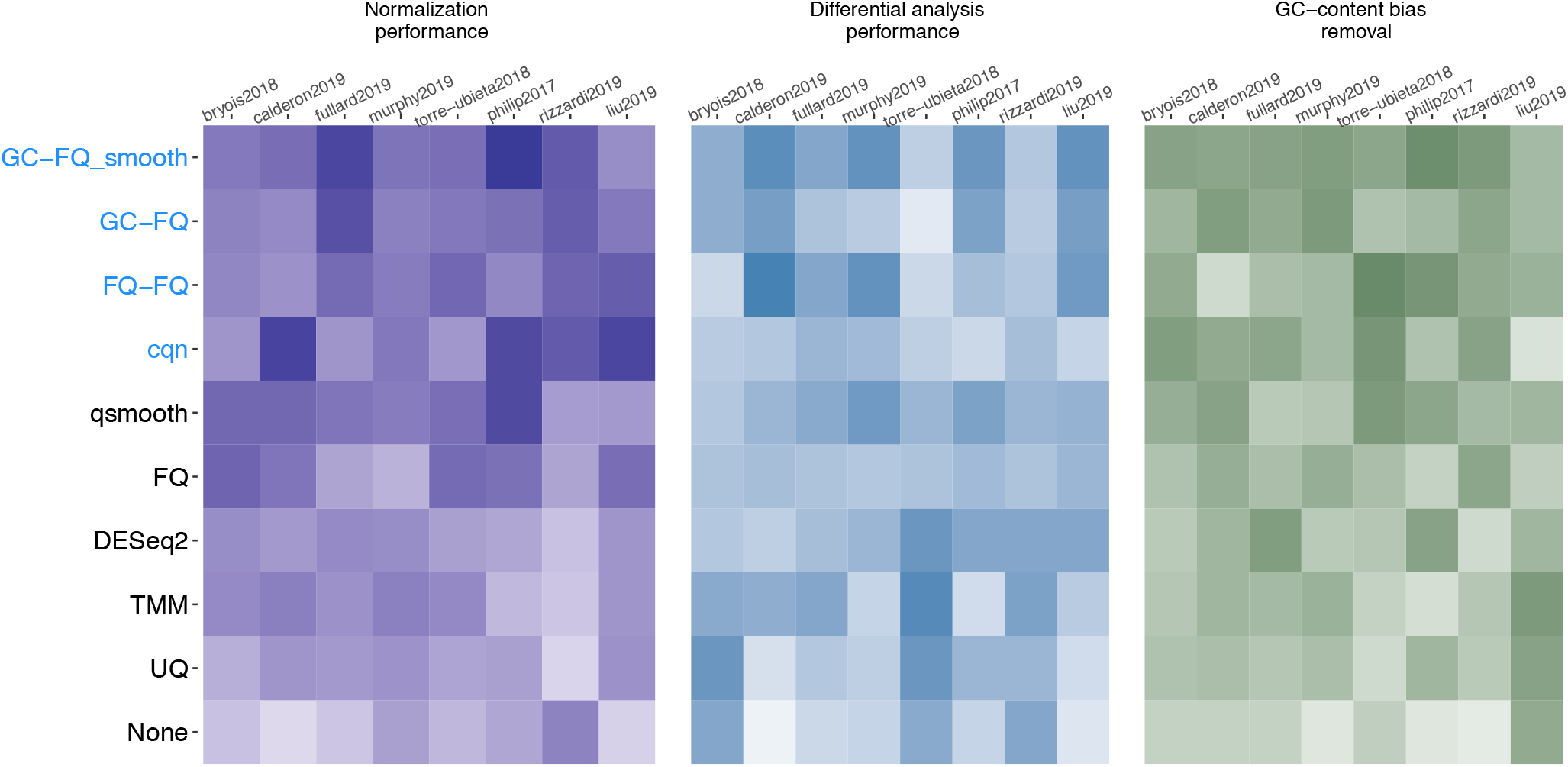
Benchmark of twelve normalization methods across eight public ATAC-seq datasets. The benchmark assesses normalization, differential accessibility performance, and GC-content bias removal, the results of which are each represented as a heatmap. The pseudo-color images, where darker color represents better performances, display matrices of average ranks (see Methods), with rows corresponding to normalization procedures and columns to datasets. Methods are ordered according to their average rank across all evaluation criteria and datasets, and their names colored based on whether they explicitly account for GC-content (blue) or not (black).

To check the robustness of these results, we also rank normalization methods for each of the three benchmarking components separately (Supplementary Figure 6). In terms of both scone normalization performance as well as GC-content effect removal, all GC-aware methods perform better than all GC-unaware methods. In terms of differential analysis performance, smooth GC-FQ is still the top performing method, followed by qsmooth and TMM normalization. GC-FQ and FQ-FQ aslo show fairly consistent good performances. Interestingly, while cqn performs well in terms of normalization performance as well as GC-content bias removal, it performs badly in differential analysis performance, being second-to-last. These results confirm that, even for benchmarking methods not explicitly using GC-content bias removal for evaluation, accounting for GC-content bias is beneficial for the normalization of ATAC-seq datasets.

Taken together, our evaluation findings show that accounting for GC-content effect is critical for normalization of ATAC-seq datasets, and, in particular, smooth GC-FQ provides good results across most datasets.

### 2.4 Case studies

In what follows, we consider the normalization of ATAC-seq datasets in greater depth using two case studies. These serve as demonstrations of how one can evaluate normalization procedures in practice using exploratory data analysis techniques. We assess each of the normalization methods that were benchmarked above, except for the basic total-count and upper-quartile normalization methods.

#### 2.4.1 Mouse Tissue Atlas

Liu et al. ^30^ presented an ATAC-seq atlas of 20 tissues in adult mice, consisting of 296, 416 peaks across 66 samples. Hierarchical clustering based on the Euclidean distance of the log-transformed normalized counts shows that normalization is essential to derive a biologically sensible clustering of the samples (Supplementary Figure 7). Without normalization, several tissues do not cluster together. The clustering is improved by using FQ normalization or global-scaling normalization methods TMM and DESeq2, but these still fail to cluster the ovary and adrenal gland tissue samples properly. By contrast, GC-aware methods cqn, FQ-FQ, and smooth GC-FQ successfully group the samples of each tissue type together, while GC-FQ misclusters one adrenal gland sample.

Next, we perform a differential accessibility analysis using the normalized counts from each normalization method as input (see Methods on how normalized counts were obtained). We model the accessibility counts using a negative binomial distribution as implemented in edgeR (or DESeq2 for DESeq2 normalization) and assess DA between heart and liver tissues. Assuming that either a small fraction of peaks are DA, or that there is symmetry in the direction of DA between the groups under comparison, log-fold-changes should be centered at zero and similarly distributed across different GC-content bins. However, log-fold-changes are biased for peaks with both low GC-content and high GC-content values for all GC-unaware normalization methods (Supplementary Figure 8). While this technical artefact is successfully removed by FQ-FQ and GC-FQ normalizations, cqn and smooth GC-FQ still suffer from substantial bias (Supplementary Figure 8). Since a high GC-content is also associated with a high accessibility count, which is in turn associated with high statistical power, we naturally expect the top DA peaks to be skewed in terms of GC-content, i.e., we expect a dominance of high GC-content values for the DA peaks. This is indeed the case for all normalization methods (Supplementary Figure 9), except TMM normalization for which the top peaks are remarkably balanced across GC-content bins. If we focus on the significant peaks at a nominal false discovery rate (FDR) threshold of 5% ^35^, most methods discover a comparable number of around 130 × 10^3^ peaks. However, cqn flags a substantially higher number of peaks, ∼ 153 × 10^3^, and therefore seems likely to return more false positives. The peaks discovered by cqn are also more balanced with respect to GC-content, as compared to other methods (Supplementary Figure 10).

#### 2.4.2 Brain Open Chromatin Atlas

Fullard et al. ^29^ published the Brain Open Chromatin Atlas (BOCA), where chromatin accessibility is measured in five human postmortem brain samples. The dataset consists of a total of 14 brain regions and two cell types (neuronal and glial/non-neuronal). These 14 brain regions can be classified into six broader regions, namely, the neocortex (NCX), primary visual cortex (PVC), amygdala (AMY), hippocampus (HIP), mediodorsal thalamus (MDT), and striatum (STR). After normalizing the counts using each normalization method (see Methods), we first assess how well each normalization method is able to recover the cell types and the major brain regions within each of these cell types by clustering the datasets using partitioning around medoids (PAM) based on the first 2 to 10 principal components. We consider PAM clustering at two resolution levels: First, we search for two clusters and check how well these correspond to the known cell types (i.e., glial and neuronal); next, we search for 12 clusters and check how well these correspond to the known regions within each cell type. We evaluate the clusterings using the adjusted Rand index (ARI) ^36,37^, comparing the derived partitions with the ground truth. Interestingly, all methods typically cluster the majority or all samples correctly according to cell type, except for cqn and no normalization (Supplementary Figure 12). However, when checking how well the different brain regions within each cell type are recovered by clustering the normalized datasets into 12 clusters (Figure 4b), for each selected number of PCs, GC-FQ, smooth GC-FQ, and FQ-FQ perform best, while cqn and no normalization perform worst. The good performance of (smooth) GC-FQ and FQ-FQ was already noticeable in the PCA plots in Supplementary Figure 11, since these are the only methods where the STR and MDT regions are clearly separated from other brain regions for the neuronal cells in 2 dimensions.

**Figure 4:**
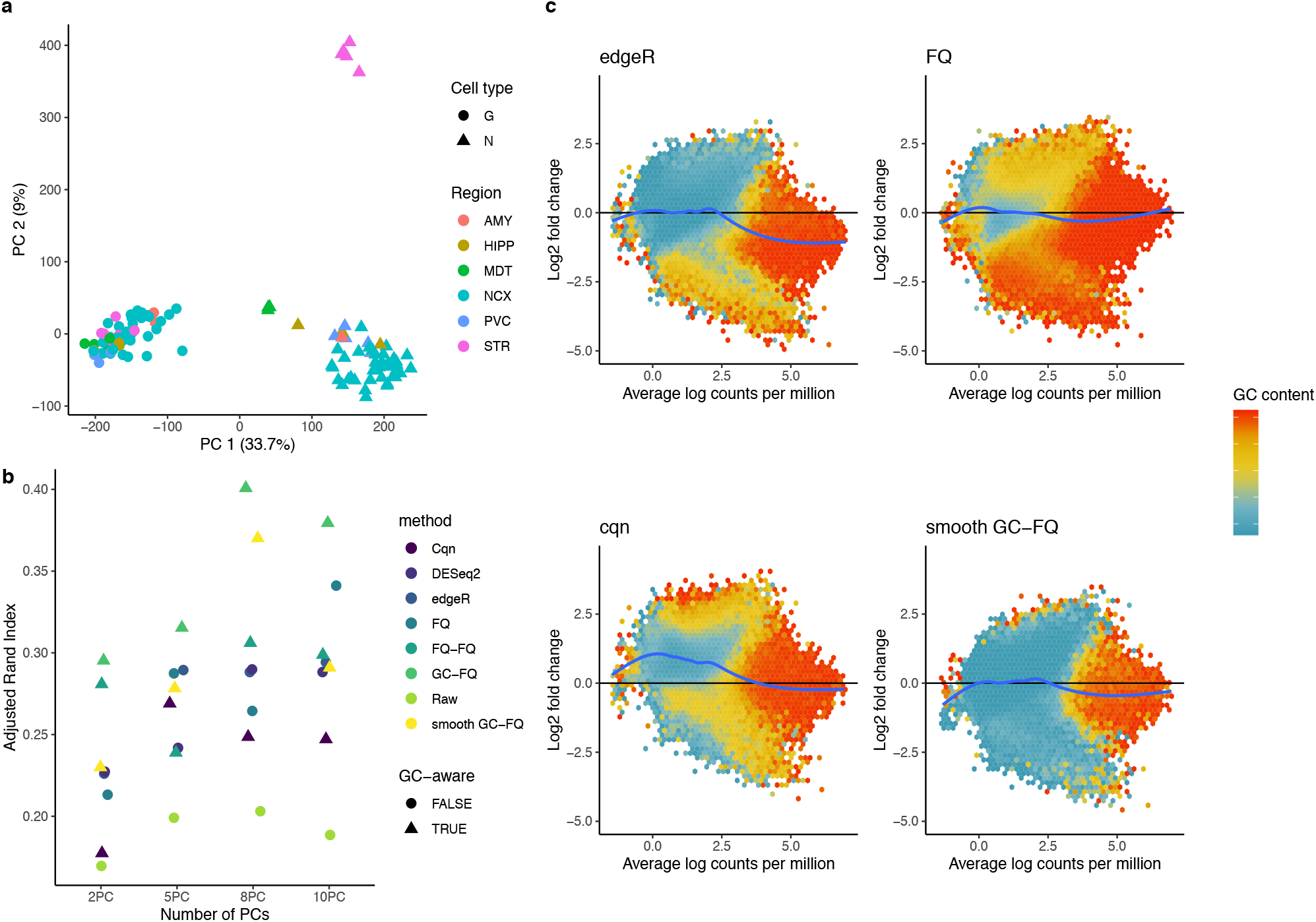
Analysis of the Brain Open Chromatin Atlas dataset. (a) PCA plot of the dataset after smooth GC-FQ normalization. The plotting symbols denote cell type, neuronal (N) and glial (G); the colors represent the six broad brain regions. (b) The samples were clustered using PAM based on a variable number of PCs (x-axis), after normalization with each of nine methods. The y-axis corresponds to the adjusted Rand index comparing the PAM clusters with the true partitioning according to brain region and cell type (12 clusters in total). Different normalizations are represented by different colors and GC-aware normalization methods are represented with triangles. GC-aware methods generally perform better, on average. (c) MD-plots for differential accessibility analysis comparing neuronal vs. non-neuronal cells. The peaks are grouped into hexagons, where the color of each hexagon denotes the average GC-content of its corresponding peaks. There is substantial GC-content bias for GC-unaware normalization methods edgeR and FQ, and similarly for all other GC-unaware methods (Supplementary Figure 13) where low GC-content is associated with high fold-changes and vice versa. The log-fold-change distribution for cqn is skewed towards positive values, also see Supplementary Figure 15. These issues are alleviated for GC-aware normalization smooth GC-FQ.

Asides from clustering, researchers often focus on discovering peaks that are differentially accessible between biological groups. Here, we use edgeR (or DESeq2 for DESeq2 normalization) to fit a negative binomial generalized linear model for each peak in each normalized dataset. For each peak, we test for differences in average accessibility between neuronal and non-neuronal cells, across all brain regions. Mean-difference plots (MD-plots) ^38^ show a prominent GC-content bias in the fold-changes for all GC-unaware normalization methods (Figure 4c and Supplementary Figure 13). Likewise, stratified violin plots of the fold-changes by GC-content show substantial bias of the fold-changes as a function of GC-content (Supplementary Figure 14). FQ-FQ and GC-FQ successfully remove GC-content effects on the fold-changes, while smooth GC-FQ removes the bias partially. Interestingly, the log-fold-changes following cqn normalization tend to be biased (Supplementary Figure 15). The peculiar results for cqn normalization are also reflected in the number of DA peaks, which is at least ∼20% higher as compared to all other methods (Supplementary Figure 16). These results emphasize the need for exploratory data analysis in order to choose an appropriate normalization method.

To assess the relevance of the discovered DA peaks, we check the genomic features and enriched gene sets associated with them, where we assign a gene to a peak if its promoter is within a 5, 000bp distance of the peak. The intersection of 134, 601 DA peaks discovered across all methods is enriched in genomic features such as exons, promoters, and 5′ UTRs, while depleted in intergenic regions, as compared to the background of all peaks (Supplementary Figure 17). The enriched biological process (BP) gene sets are highly relevant, including *neurogenesis* and *nervous system development*, among others (Supplementary Table 1). The DA results also allow us to assess whether accounting for GC-content effects can aid biological interpretation. We therefore interpret the set of 9, 866 peaks discovered by FQ-FQ, smooth GC-FQ, and GC-FQ, while not by their GC-unaware counterpart, FQ normalization. These peaks are enriched in genomic features such as promoters, 5′ UTRs, and exons (Supplementary Figure 17). While no biological process gene sets are significantly enriched at a 5% FDR level, the top gene sets are still relevant, including *regulation of synapse structure or activity* and *synapse organization* (Supplementary Table 2). We also further investigate the peaks uniquely discovered by cqn. These peaks are enriched in intergenic regions

(Supplementary Figure 17), which supports our intuition that these are likely false positive peaks. While again no enriched gene sets are found at a nominal 5% FDR level, in this case the top gene sets are not relevant to the experiment, mostly involving gene sets on the kidney and eye (Supplementary Table 3), reinforcing the hypothesis that these could be false positives.

Taken together, these results again suggest that GC-content normalization is crucial for the analysis of ATAC-seq data, improving downstream analyses and biological interpretation. Exploratory data analysis is essential for evaluating and guiding the choice of effective normalization and removal of technical GC-content effects.

## 3 Discussion

The evaluations in this manuscript highlight the importance of accounting for GC-content effects in ATAC-seq datasets. Because of the sample-specificity of GC-content effects, failing to adjust for GC-content using an appropriate normalization method can bias downstream analyses such as clustering and differential accessibility analysis. We have proposed GC-aware normalization procedures and benchmarked these against state-of-the-art procedures using eight public ATAC-seq datasets. While GC-aware procedures perform better than GC-unaware procedures, none uniformly outperforms all others, although smooth GC-FQ generally performs well on average. The choice of an appropriate normalization procedure is dataset-specific and exploratory data analysis is essential to guide this choice.

Similar GC-content effects have also been noted in DNA-seq ^39^, RNA-seq ^11,21^, and ChIP-seq ^9^, amongst others. For ChIP-seq datasets, Teng and Irizarry ^9^ recently developed a negative binomial mixture model to correct for GC-content effects in both background and binding signal regions at the peak-calling stage, by accounting for GC-content in the abundance estimation for a particular genomic region. While their method has been evaluated using ChIP-seq data, it may also be useful for ATAC-seq data. However, none of the publicly-available and processed datasets we have looked into account for GC-content effects during the peak-calling stage.

While in this manuscript we have focused on correcting GC-content bias at the level of called peaks, other approaches are possible. For example, Benjamini and Speed ^39^ argue that it is the GC-content of the full DNA fragment (vs. only the sequenced read) that most influences read counts. A comparison of the peak-level normalization approaches discussed here with fragment-level approaches would be an interesting avenue for further research on how to best correct for GC-content effects.

Our work has focused on normalization of bulk ATAC-seq datasets. While FQ-based normalization procedures were found to perform favorably in this setting, it remains to be seen whether they perform equally well on single-cell ATAC-seq (scATAC-seq) datasets. The sparsity associated with scATAC-seq data suggests that their application could be limited and alternative normalization procedures may be needed.

## 4 Methods

### 4.1 Datasets

Philip et al. ^17^ study CD8 T-cell dysfunction in acutely infected and chronic tumoral tissue, over several time points. We only focus on the mouse samples and consider time and treatment as the biological variables of interest. We did not find metadata on quality control (QC) or batch variables, so we do not use any in the scone evaluation. The count matrix corresponds to 75, 689 peaks for 41 samples and was downloaded from the Gene Expression Omnibus (GEO) with accession number GSE89308.

Bryois et al. ^28^ study the adult human prefrontal cortex brain. We remove samples that are not schizophrenic or control samples, leaving a total of 272 samples, consisting of 135 individuals with schizophrenia and 137 controls. We consider the disease status as the biological variable of interest. In the evaluation, we use the 32 QC variables that were available in the metadata, along with the top 10 principal components derived from the patients’ genotypes. The sequencing index in the metadata is used as batch variable. The count matrix corresponds to 118, 152 peaks and was obtained through personal communication with the authors. It was not relevant to correct for the width of the peaks using cqn in this dataset, since all peaks have a length of 301bp.

de la Torre-Ubieta et al. ^14^ study human cortical neurogenesis in the germinal zone and cortical plate of the developing cerebral cortex. Samples were derived from three individual donors and each donor was handled and processed separately, so we treat each donor as a batch and the brain region as the biological variable of interest. The count matrix corresponds of 62, 005 peaks across 19 samples and was downloaded from the GEO with accession number GSE95023. Note that the replication in this dataset is technical, i.e., consists of samples from the same human donor.

Calderon et al. ^23^ study a repertoire of 32 immune cell types under resting and activated conditions in humans.

The metadata include three QC variables (number of peaks called, number of sequenced reads, and transcription start site enrichment for each sample), which we use in the scone evaluation. Most donors are processed and sequenced separately, and therefore each donor represents a different batch. However, some samples underwent a second round of sequencing and the accessibility counts from these two sequencing rounds were summed, so we also treat these additionally sequenced samples as a separate batch. The biological variables of interest are cell type and treatment. The count matrix corresponds to 829, 942 peaks across 175 samples and was downloaded from the GEO with accession number GSE118189. This dataset was filtered to retain peaks with at least 2 counts per million in at least 10 samples, reducing the dataset to 203, 448 peaks.

Murphy et al. ^31^ study photoreceptors and bipolar cells in the mouse retina. The dataset combines two experiments and we define each experiment as a batch. We consider the cell type as the biological variable of interest. The count matrix corresponds to 110, 715 peaks across 12 samples and was downloaded from the GEO with accession number GSE131625. It was not relevant to correct for the width of the peaks using cqn in this dataset, since all peaks have a length of 201bp.

Rizzardi et al. ^16^ study neuronal and non-neuronal cell populations in the prefrontal cortex and nucleus accumbens in humans. We consider the combination of brain region and cell type as the biological variable of interest. We define a batch variable as the combination of the donor, flow cytometry run, and sequencing date variables. The count matrix corresponds to 961, 916 peaks across 22 samples and was downloaded from the GEO with accession number GSE96614.

#### Brain Open Chromatin Atlas (case study)

Fullard et al. ^29^ developed a human brain atlas of neuronal and non-neuronal cells across 14 distinct brain regions from 5 human donors. We define a batch variable as the flow cytometry date. Note that while the sequencing date is nested within the flow cytometry date, there are other variables in the metadata that might also be considered to define batches, e.g., PCR date. A total of 49 variables corresponding to potential technical effects were included as QC measures. The biological variable of interest is defined as the combination of cell type and brain region. The count matrix corresponds to 300, 444 peaks across 115 samples and was downloaded from the Brain Open Chromatin Atlas (BOCA) website, at https://bendlj01.u.hpc.mssm.edu/multireg/.

#### Mouse Tissue Atlas (case study)

Liu et al. ^30^ created an ATAC-seq atlas of mouse tissues, spanning a total of 20 tissues for both male and female mice. We use the “Slide lane of sequencer” variable recorded in the metadata as batch variable. Four variables (mitochondrial reads, usable reads, transcription start site (TSS) enrichment, and number of reproducible peaks, as identified using the irreproducible discovery rate (IDR) ^40^) are used as QC measures. The combination of gender and tissue type is used as the biological variable of interest. The count matrix corresponds to 296, 574 peaks across 79 samples and was downloaded from Figshare at https://doi.org/10.6084/m9.figshare.c.4436264.v1.

#### 4.2 GC-content retrieval

For each dataset, we use the Bioconductor R package Biostrings to retrieve the GC-content of every peak region, using the reference genome of the relevant organism. Table 1 provides the genome version used for each dataset.

**Table 1:** Genome version used for each dataset.

| Dataset | Organism | Genome |
| --- | --- | --- |
| Philip et al.<sup>17</sup> | Mouse | GRCm38 |
| Bryois et al.<sup>28</sup> | Human | GRCh37.75 |
| Calderon et al.<sup>23</sup> | Human | GRCh37.75 |
| de la Torre-Ubieta et al.<sup>14</sup> | Human | GRCh37.75 |
| Murphy et al.<sup>31</sup> | Mouse | GRCm38 |
| Rizzardi et al.<sup>16</sup> | Human | GRCh37.75 |
| Fullard et al.<sup>29</sup> | Human | GRCh37.75 |
| Liu et al.<sup>30</sup> | Mouse | GRCm37.67 |

### 4.3 Normalization procedures

Let *Y*_*ji*_ denote the accessibility count for peak *j* = 1,…, *J* in sample *i* = 1,…, *n*. The evaluated normalization procedures can be summarized as follows.

#### No normalization

The raw counts are used for analysis.

#### Total-count (TC) normalization

Each count is divided by the total library size, *N*_*i*_ = Σ_*j*_ *Y*_*ji*_, for its corresponding sample.

#### Upper-quartile (UQ) normalization

Each count is divided by the upper-quartile (i.e., 75th percentile) of the counts for its corresponding sample. UQ can be beneficial over TC normalization, as the latter can be affected by a few very high counts that dominate the total library size *N*_*i*_.

#### Trimmed mean of *M* -values (TMM) normalization

TMM is a global-scaling normalization procedure that was originally proposed by Robinson and Oshlack ^19^. As the name suggests, it is based on a trimmed mean of fold-changes (*M* -values) as the scaling factor. A trimmed mean is an average after removing a set of “extreme” values. Specifically, TMM calculates a normalization factor 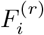 for each sample *i* as compared to a reference sample *r*,

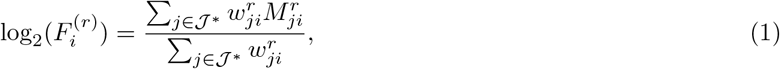

where 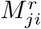 represents the log_2_-fold-change of the accessibility fraction as compared to a reference sample *r*, i.e.,

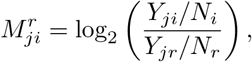

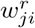 represents a weight calculated as

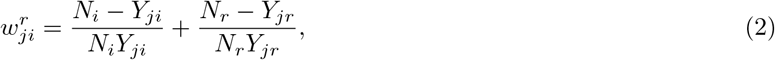

and 𝒥* represents the set of peaks after trimming those with the most extreme values.

The procedure only takes peaks into account where both *Y*_*ji*_ *>* 0 and *Y*_*jr*_ *>* 0. By default, TMM trims peaks with the 30% most extreme *M* -values and 5% most extreme average accessibility, and chooses as reference *r* the sample whose upper-quartile is closest to the across-sample average upper-quartile. The normalized counts are then given by 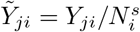, where

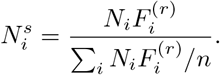

#### DESeq2 normalization

The median-of-ratios method is used in DESeq2^20^. It assumes that the expected value *µ*_*ji*_ = *E*(*Y*_*ji*_) is proportional to the true accessibility of the peak, *q*_*ji*_, scaled by a normalization factor *s*_*i*_ for each sample,

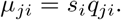

The normalization factor *s*_*i*_ is then estimated using the median-of-ratios method compared to a synthetic reference sample *r* defined based on geometric means of counts across samples

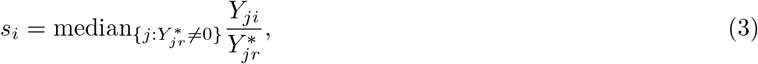

with

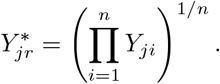

From this, we calculate the normalized count as 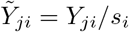.

#### Full-quantile (FQ) normalization

In full-quantile normalization ^41^, the samples are forced to each have a distribution identical to the distribution of the median/average of the quantiles across samples. In practice, we implement full-quantile normalization using the following procedure

1. given a data matrix **Y**_*J*×*n*_ for *J* peaks (rows) and *n* samples (columns),
2. sort each column to get **Y**^*S*^,
3. replace all elements of each row by the median (or average) for that row,
4. obtain the normalized counts 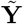 by re-arranging (i.e., unsorting) each column.

#### Smooth-quantile (SQ) normalization (qsmooth)

Full-quantile normalization assumes that the read count distribution has a similar shape for each sample and that the observed variability in global distributional properties corresponds to technical effects. However, this may not always be the case in practice. To tackle this, Hicks et al. ^42^ developed smooth-quantile normalization, a variant of full-quantile normalization that is able to deal with datasets where there are large global differences between biological conditions of interest. It provides a balance between (a) full-quantile normalization between samples of each condition separately, and (b) full-quantile normalization on the full dataset. This balance is struck by calculating data-driven weights for each quantile, that specify which of the two normalization options is more appropriate. The weights are estimated in a smooth way across the quantiles, by contrasting the within-condition with the between-condition variability for each quantile. If the within-condition variability is significantly smaller than the between-condition variability, then the weights will favor normalization for each condition separately.

#### Within-and-between-sample full-quantile (FQ-FQ) normalization

The FQ-FQ method, implemented in the EDASeq package ^10^, accounts for GC-content effects by performing two rounds of full-quantile normalization. First, the features of each sample are grouped into (by default, 10) GC-content bins and full-quantile normalization is performed across bins within each sample (referred to as ‘within-lane normalization’). Next, the data are normalized using full-quantile normalization across all samples.

#### Conditional-quantile normalization (cqn)

The cqn method ^43^ starts by assuming a Poisson model for the accessibility counts *Y*_*ji*_. Median regression is used to model, for each sample, the log-transformed accessibility count as a smooth function of GC-content as well as peak width, focusing on peaks with high average count (above 50 by default). Note that for the datasets from Bryois et al. ^28^, Murphy et al. ^31^, and Rizzardi et al. ^16^ all peaks have the same width and hence there is no peak width normalization. Next, subset quantile normalization ^44^ is performed on the residuals of that model (i.e., on the counts adjusted for GC-content) for between-sample normalization. The method could intuitively be thought of as full-quantile normalization after removing a smoothed sample-specific GC-content effect. Normalized counts are calculated as recommended in the cqn vignette, i.e.,

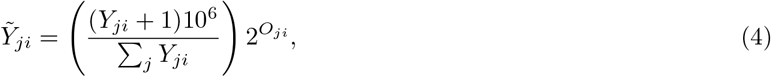

with *O*_*ji*_ the normalization offset estimated by cqn, which is on the log_2_ scale.

#### GC-full-quantile (GC-FQ) normalization

GC-FQ is similar to FQ-FQ, but relies on the observation that, in ATAC-seq, read count distributions are often more comparable between samples within a GC-content bin, than between GC-content bins within a sample (Figure 2). It therefore applies between-sample FQ normalization for each GC-content bin separately, with 50 bins by default.

#### Smooth GC-FQ normalization

smooth GC-FQ is a variant of GC-FQ that applies smooth-quantile normalization across samples within each GC-content bin. Like GC-FQ, it uses 50 bins by default.

### 4.4 Benchmarking

#### 4.4.1 Normalization performance: scone benchmark

We use the Bioconductor R package scone ^33^ to implement and evaluate different normalization procedures. The first step in the scone workflow is to normalize the data using all normalization methods of interest. The normalized data are then used to calculate a range of evaluation measures (see Supplementary Methods for a description and evaluation of each of these). Since some measures tend to be biased towards particular normalization methods, we rely on a subset, selected based on our evaluation as described in Supplementary Results. As part of the evaluation, the log-count principal components of the normalized data are correlated to factors of unwanted variation as well as quality control variables (if available). The factors of unwanted variation are inferred using RUVSeq ^45^, based on negative control features which we here set to be the peaks within known housekeeping genes.

#### 4.4.2 Differential accessibility analysis performance in a synthetic null and signal scenario

Our approach to evaluate the impact of normalization on DA analysis is two-fold: First, we perform synthetic null comparisons for each real dataset; second, we generate synthetic signal datasets by simulating DA peaks from each real dataset (see Methods, Datasets for a descriptions of each dataset).

##### Synthetic null scenario

In the null scenario, for each dataset, we create a two-group mock variable so that we expect no systematic differences between the groups. Specifically, for each dataset, we perform stratified random sampling, where the samples for each biological condition are randomly split into two approximately equally-sized groups (e.g., for a biological condition with 4 samples, there are 2 samples per group, and for a biological condition with 3 samples, one group comprises 1 sample and the other 2 samples). For each dataset, following the assignment of samples to groups, we evaluate the performance of normalization methods based on a differential accessibility analysis using this mock variable.

##### Sythetic signal scenario

Additionally, we also evaluate the performance of normalization and DA analysis methods on synthetic signal datasets created from each of the real datasest and where 10% of all peaks are DA.

Each synthetic dataset comprises 12 samples (based on 6 randomly selected samples from each group in the mock variable). For each selected sample *i*, we calculate its accessibility fraction for each peak *j* as

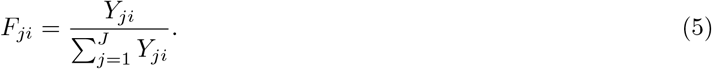

A random subset comprising 10% of all peaks is simulated to be differentially accessible, with equal up-/downregulation between groups, via independent binary random variables *S*_*j*_, equal to either −1 or 1, each with 1*/*2 probability. The *S*_*j*_’s define the group *g* ∈ {1, 2} of samples for which the accessibility fractions will be altered as follows

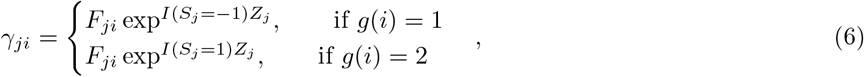

where *g* ∈ {1, 2} denotes the group to which sample *i* belongs and *Z*_*j*_ are independent Gaussian random variables with mean 0.8 and standard deviation 0.1. That is, the log-fold-change in accessibility between groups 2 and 1 is *S*_*j*_*Z*_*j*_. The choice of the mean and standard deviation for the log-fold-changes corresponds to fold-changes being on average 2.25, with a minimum of ∼ 1.5 and a max of ∼ 3.5, a reasonable scenario. Sequencing depths 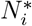 are simulated from a uniform distribution

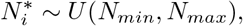

where *N*_*min*_ = min_*i*_ Σ_*j*_ *Y*_*ji*_ and *N*_*max*_ = max_*i*_ Σ_*j*_ *Y*_*ji*_ denote, respectively, the minimum and maximum library sizes across all samples in the dataset. Given *γ*_*ji*_ and 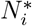, accessibility counts 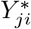 are then simulated using a Multinomial distribution

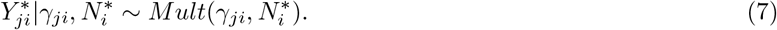

For each combination of sample size and DA signal strength, we evaluate 14 simulated datasets (a greater number of simulations was not feasible due to local memory limitations). Differential accessibility analysis is performed using edgeR for all normalization methods, except for DESeq2 normalization where we rely on the native DESeq2 pipeline.

For RUVg normalization, we use peaks overlapping with housekeeping genes as negative control features, as in the scone evaluation, excluding features that were simulated to contain signal between the groups.

For each simulated dataset, we calculate the true positive rate (TPR) and false discovery rate (FDR), defined as

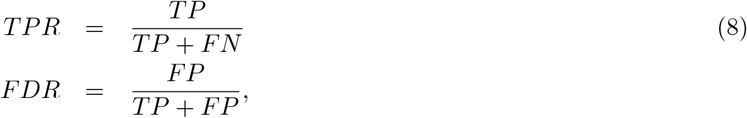

where *FN, FP*, and *TP* denote, respectively, the numbers of false negatives, false positives, and true positives. Method performance is visualized using FDR-TPR curves, constructed by calculating, for each of the 14 simulated datasets, FDR and TPR ratios by sequentially moving from the most to the least significant DA peak and then averaging the FDR and TPR over the 14 simulations.

#### 4.4.3 Method ranking

Each normalization procedure is assigned a score for each evaluation criterion, constructed such that a high score corresponds to a good performance of the normalization procedure with respect to that evaluation criterion. Since scores are not directly comparable between criteria, we first rank the normalization methods for each evaluation criterion separately, where a high rank reflects good normalization. As a summary for each normalization procedure, we average the ranks across evaluation criteria.

### 4.5 Case studies

#### 4.5.1 Mouse Tissue Atlas

The raw count matrix from Liu et al. ^30^ is obtained as described in the Datasets section and is normalized using each of the twelve evaluated normalization procedures, with no peaks filtered out. Euclidean distances between samples are calculated on the log-normalized counts, adding an offset of 1 to avoid taking the log of zero. Hierarchical clustering trees are derived using complete linkage. Differential accessibility analysis is performed using edgeR for all normalization methods, except for DESeq2 normalization where we rely on the native DESeq2 pipeline. Normalized counts are used directly as input for FQ, FQ-FQ, and (smooth) GC-FQ normalization, while normalization offsets are used for TMM and cqn. For RUVSeq normalization, we incorporate 4 inferred factors of unwanted variation in the design matrix.

#### 4.5.2 Brain Open Chromatin Atlas

The raw count matrix from Fullard et al. ^29^ is obtained as described in the Datasets section. We do not filter out any peaks. PCA and hierarchical trees are based on the log-normalized counts, adding an offset of 1 to avoid taking the log of zero. Hierarchical trees use the Euclidean distance between samples and are constructed using complete linkage. DA analysis is performed as described in the Mouse Tissue Atlas case study. The contrast matrix is defined for comparing the average expression of neuronal vs. non-neuronal samples.

## 5 Data and code availability

A GitHub repository containing all code for the analyses can be found at https://github.com/koenvandenberge/ `bulkATACGC`, which also contains a link to Zenodo to download the data used for these analyses. All datasets used in this study are publicly-available, either online or by contacting the original authors, as described in the Methods section.

## 6 Acknowledgements

The authors thank Michael Love for his input on scaling normalization methods edgeR and DESeq2, and Ameek Bindra, who contributed to this project through the Undergraduate Research Apprenticeship Program (URAP) of the Department of Statistics at the University of California, Berkeley.

KVdB is a postdoctoral fellow of the Belgian American Educational Foundation (BAEF) and is supported by the Research Foundation Flanders (FWO) grants 1246220N and G062219N. H-J C was supported by a postdoctoral research grant from the UC Berkeley Siebel Stem Cell Center. SD and JN are supported by the National Institutes of Health, grant R01 DC007235.

