## Supplementary Material for "Normalization benchmark of ATAC-seq datasets shows the importance of accounting for GC-content effects"

### 1 Supplementary Methods

#### 1.1 Defining scone evaluation measures

##### 1.1.1 Default scone evaluation measures

By default, scone uses a range of evaluation measures to assess normalization. The relevant measures for this work can be divided into three categories, where we use the definitions from the scone paper[33].

**Clustering properties.**

- *BIO_SIL*: Group the samples according to the value of a categorical covariate of interest (e.g., known cell type, genotype) and compute the average silhouette width for the resulting clustering.
- *BATCH_SIL*: Group the samples according to the value of a nuisance categorical covariate (e.g., batch) and compute the average silhouette width for the resulting clustering.
- *PAM_SIL*: Cluster the samples using partitioning around medoids (PAM) for a range of user-supplied numbers of clusters and compute the maximum average silhouette width for these clusterings.

**Association with control genes and QC measures.**

- *EXP_QC_COR*: The weighted coefficient of determination (see Cole et al.[33] for details) for the regression of log-count principal components on all principal components of user-supplied QC measures.
- *EXP_UV_COR*: The weighted coefficient of determination (see Cole et al.[33] for details) for the regression of log-count principal components on factors of unwanted variation (default 3) derived from negative control genes.

**Global distributional properties.**

- *RLE_MED*: Mean squared median relative log-expression (RLE).
- *RLE_IQR*: Variance of inter-quartile range (IQR) of RLE.

##### 1.1.2 GC-content bias evaluation measures

To assess GC-content bias after normalization, we define two measures based on relative log-expression values[34] across GC-content bins, which are inspired by the RLE_MED and RLE_IQR measures already implemented in scone.

Let $Y_{ji}$ denote the accessibility measure for peak $j$ in sample $i$ and $L_{ji} = \log(Y_{ji} + 1)$. Then, the RLE is defined as

$$r_{ji} = L_{ji} - \bar{L}_{j.},$$

where $\bar{L}_{j.}$ denotes the median of $L_{ji}$ across all samples $i$. For a set of peaks within a GC-content bin $b$, we can define a measure of GC-content bias as the mean squared median RLE[33], i.e., as the average squared deviation of the median RLE from zero,

$$d_b = \frac{1}{n} \sum_{i=1}^{n} \bar{r}_{.ib}^2,$$

where $\bar{r}_{.ib}$ is the median RLE value for peaks in bin $b$ for sample $i$. A small value of $d_b$ generally corresponds to a good normalization of the data, since the median RLE is close to zero. However, if sample-specific GC-content effects exist, then the normalization may only be working for certain ranges of GC-content. We therefore assess whether the mean squared median RLE varies with GC-content by computing its variance across GC-content bins,

$$RLE\_MED_{GC} = \frac{1}{B-1} \sum_{b=1}^{B} (d_b - \bar{d})^2, \qquad (9)$$

where $\bar{d} = \sum_b d_b/B$ is the average of $d_b$ across GC-content bins. In the evaluation, we let the number of bins $B$ depend on the total number of peaks in a given dataset by constructing bins containing around $4,000$ peaks each.

A similar measure is calculated based on the variance of the interquartile range of the RLE measures for each bin,

$$v_b = \frac{1}{n-1}\sum_{i=1}^{n}(q_{ib} - \bar{q}_b)^2,$$

where $q_{ib}$ is the interquartile range of the RLE values for peaks in bin $b$ for sample $i$ and $\bar{q}_b$ its average across all samples. Using a similar reasoning as above, we then evaluate the variance of $v_b$ across different GC-content bins

$$RLE\_IQR_{GC} = \frac{1}{B-1}\sum_{b=1}^{B}(v_b - \bar{v})^2, \tag{10}$$

where $\bar{v}$ is the average of $v_b$ across all bins.

#### 1.2 Differential accessibility analysis performance measures

In addition to evaluating normalization performance using scone, we also use each dataset in a differential accessibility (DA) performance evaluation, see Methods for details. The performance evaluation relies on DA analysis results on both a mock comparison of the real dataset, as well as a simulated dataset based on each real dataset. We define the following set of measures.

**Mock evaluation.**

- *False positive rate*: The false positive rate at a nominal 5% significance level for each peak. In this null setting, a good performing method is expected to have 5% of its $p$-values less than or equal to 0.05.
- *p-value uniformity*: The Hellinger distance between the observed $p$-value distribution and a uniform $p$-value distribution. A good performing method is expected to have a small distance.
- *p-value uniformity as function of GC-content*: The variability Hellinger distance between the observed $p$-value distribution and a uniform $p$-value distribution across GC-content bins. A good performing method is expected to have no GC-content bias in terms of $p$-value uniformity.

**Simulated datasets with signal.**

- *Area under receiver operator characteristic curve* (AUROC): The area under the receiver operator characteristic (TPR-FPR) curve. A good method is expected to have a high value of AUROC, i.e., is capable of identifying the truly DA peaks without simultaneously calling too many false positives.
- *GC-content distribution of DA peaks*: Distance in empirical cumulative distribution functions between the observed GC-content distribution of called DA peaks and the GC-content distribution of truly DA peaks. This distance is calculated over a grid of 100 points as the sum of the absolute differences between the empirical cumulative distribution functions of the true GC-content distribution and the observed GC-content distribution of called DA peaks. A good performing method should have a small distance.

### 2 Supplementary Results

#### 2.1 Selecting scone evaluation measures

##### 2.1.1 Selection of evaluation measures

All scone evaluation measures are described in Supplementary Methods. Since some measures are biased in favor of particular normalization procedures, here we investigate which subset of measures would be suitable to use. To select evaluation measures, we investigate the relationship of the scores with all normalization methods under three scenarios.

1. A mock benchmark using simulated data without GC-content effects: Here, we simulate data corresponding to $10,000$ peaks and 16 samples from a negative binomial distribution for each feature, using the makeExampleDESeqDataSet function in DESeq2[20]. Each feature is assigned a random GC-content value, and there is no normalization required asides from a correction for sequencing depth. Since no normalization for technical variables is necessary, this scenario allows us to check if some evaluation measures favor particular normalization methods.

2. A benchmark on all real datasets, permuting GC-content: Here, all real datasets are used, but GC-content is permuted across features, eliminating any GC-content bias. This scenario allows us to evaluate additional measures and confirm the results from the mock benchmark using real data.

3. A benchmark on all real datasets: Here, all real datasets are used. This scenario allows us to confirm whether the RLE measures assessing GC-content bias removal are useful. Indeed, since we can expect GC-content bias in this evaluation, but not in the previous one, measures that do not change between evaluations will fail to pick up GC-content bias.

**Mock benchmark.** Based on the mock benchmark, we observe that no normalization method is favored over others for the *BIO_SIL*, *PAM_SIL*, and *EXP_UV_COR* measures (Supplementary Results Figure 1). *RLE_MED* favors FQ, FQ-FQ, and GC-FQ, while *RLE_IQR* favors smooth GC-FQ and qsmooth. A similar pattern is observed for GC-aware versions of the procedures, which is good since there is no actual GC-content bias in the mock benchmark. This result makes clear that RLE measures favor methods that heavily rely on full-quantile normalization, and should therefore be avoided.

**Real data, permuted GC-content.** The real data benchmark with permuted GC-content allows us to check additional measures such as *EXP_QC_COR* and *EXP_UV_COR*, on real data where we do not expect GC-content bias. This analysis largely recapitulates the observations we made for the mock comparison in that RLE measures favor procedures that use full-quantile normalization (Supplementary Results Figure 2). As in Supplementary Results Figure 1, the GC-aware versions of the RLE measures are similar to their GC-unaware counterparts, which is again a good sign since we do not expect GC-content effects in this evaluation. In addition, measures *EXP_QC_COR* and *EXP_UV_COR* are relevant measures to be able to discriminate between normalization methods.

**Real data.** The results on the real data benchmark (Supplementary Results Figure 3) are similar to those when GC-content is permuted for all measures except $RLE\_MED_{GC}$. $RLE\_MED_{GC}$, a measure that assesses GC-content removal, is drastically different, which is encouraging, since in this scenario it is expected that GC-content bias is a major contributor to technical variability. However, the fact that $RLE\_IQR_{GC}$ is similar to *RLE_IQR* and the results when GC-content is permuted, suggests it is not a useful measure to assess removal of GC-content bias effects.

Based on these evaluations, we have selected *BIO_SIL*, *BATCH_SIL*, *PAM_SIL*, *EXP_QC_COR*, *EXP_UV_COR*, and $RLE\_MED_{GC}$ as our measures of choice in the scone benchmark. We do not use *RLE_MED* and *RLE_IQR*, since these are biased towards methods based on FQ-normalization. We also do not use $RLE\_IQR_{GC}$, since it seems unsuccessful at evaluating the removal of GC-content effects, possibly due to the mean-variance relationship of count data and the positive association of mean accessibility with GC-content, naturally causing the variance of IQR across samples to increase for bins containing more accessible peaks.

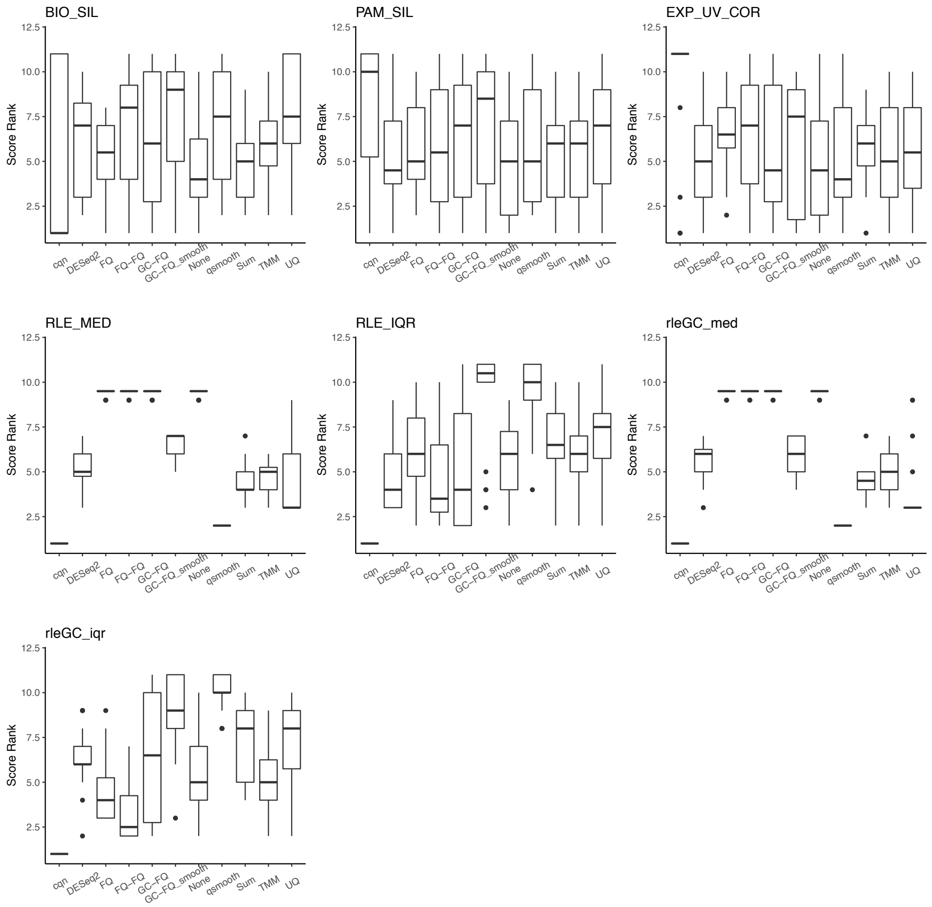


Supplementary Results Figure 1: *scone score ranks for the mock simulation benchmark.* Different panels correspond to different evaluation measures, and normalization methods are defined on the x-axis. A higher score rank means a better normalization with respect to a particular measure.

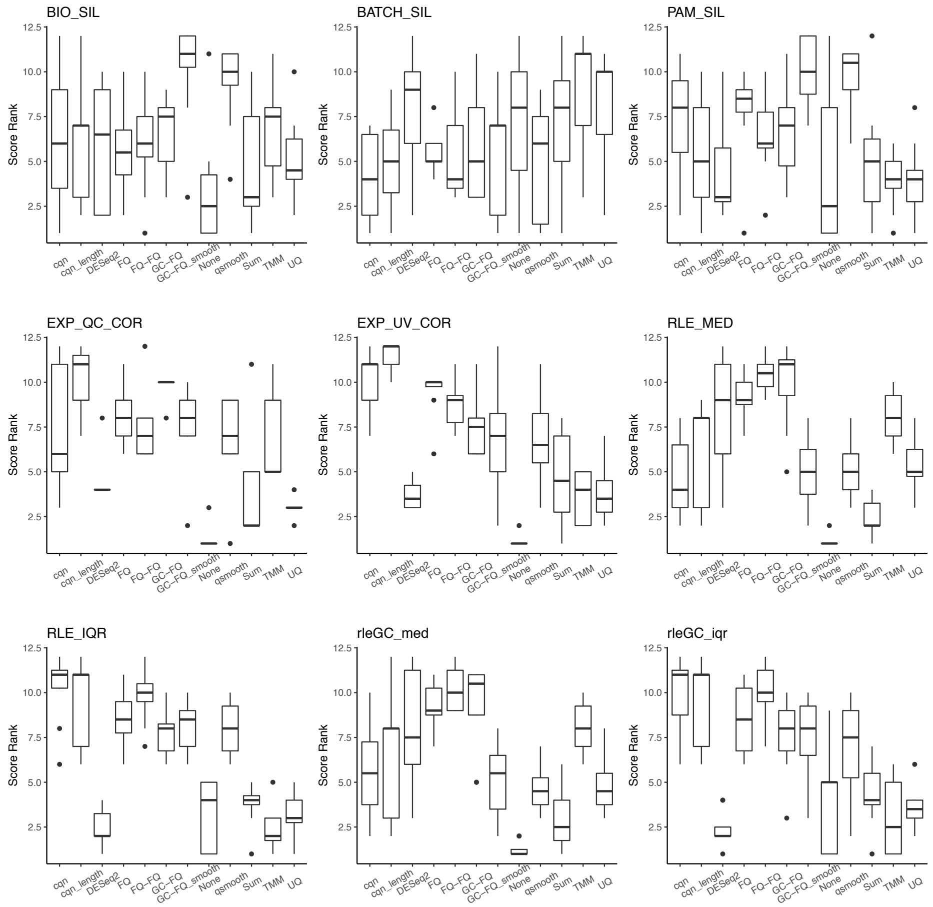


Supplementary Results Figure 2: *scone score ranks for the real data benchmark with permuted GC-content.* Different panels correspond to different evaluation measures, and normalization methods are defined on the x-axis. A higher score rank means a better normalization with respect to a particular measure.

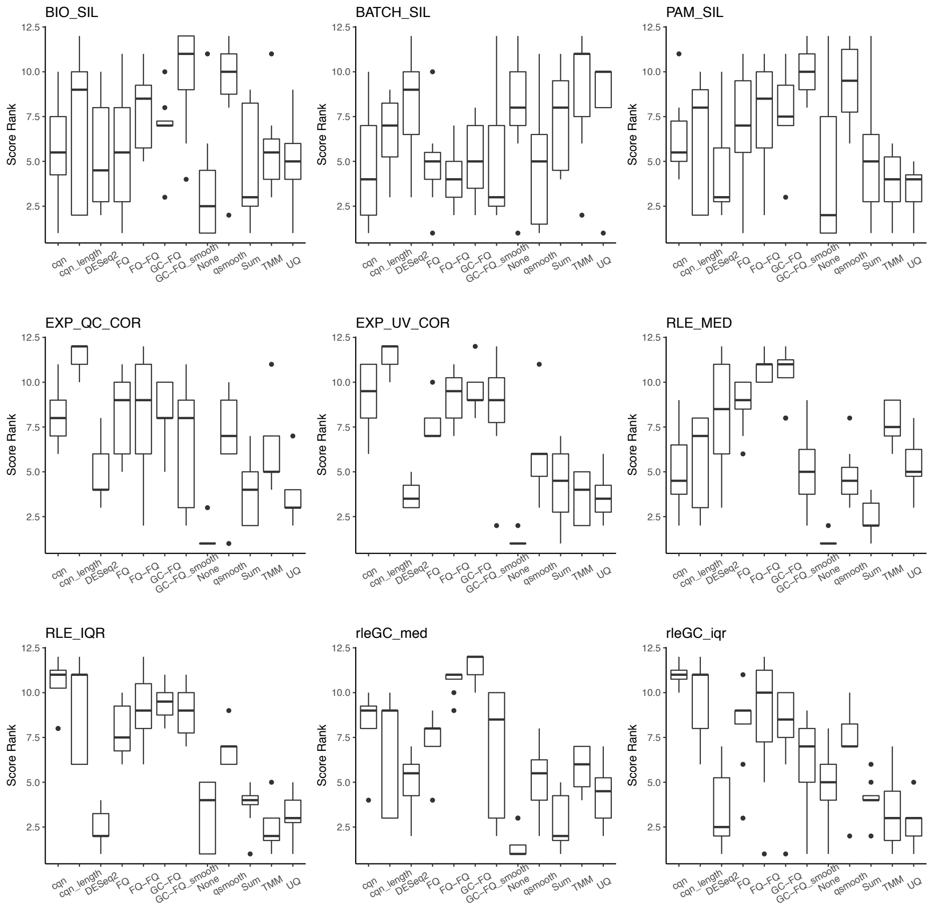


Supplementary Results Figure 3: *scone score ranks for the real data benchmark.* Different panels correspond to different evaluation measures, and normalization methods are defined on the x-axis. A higher score rank means a better normalization with respect to a particular measure.

### 3 Supplementary Figures

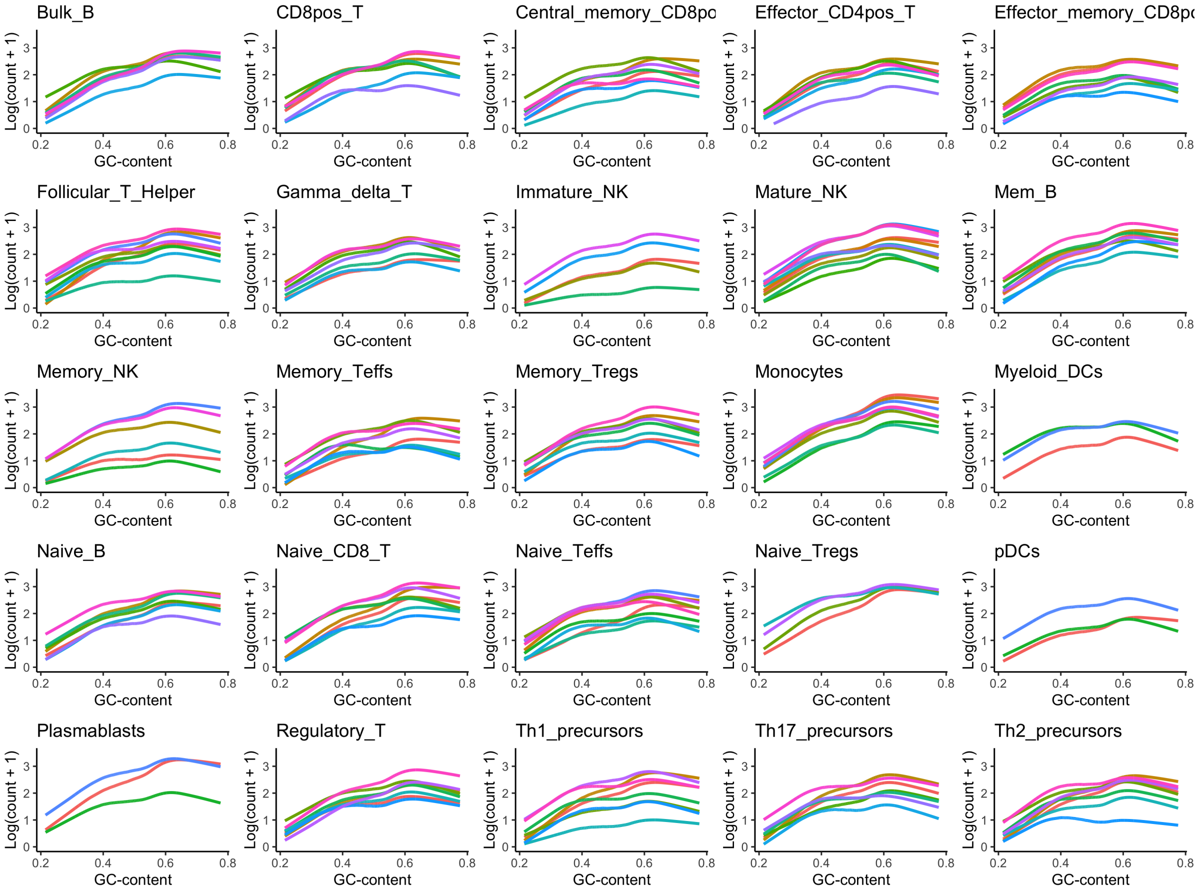


Supplementary Figure 1: *Lowess fits for the log accessibility count as a function of GC-content for the dataset from Calderon et al.*[23].

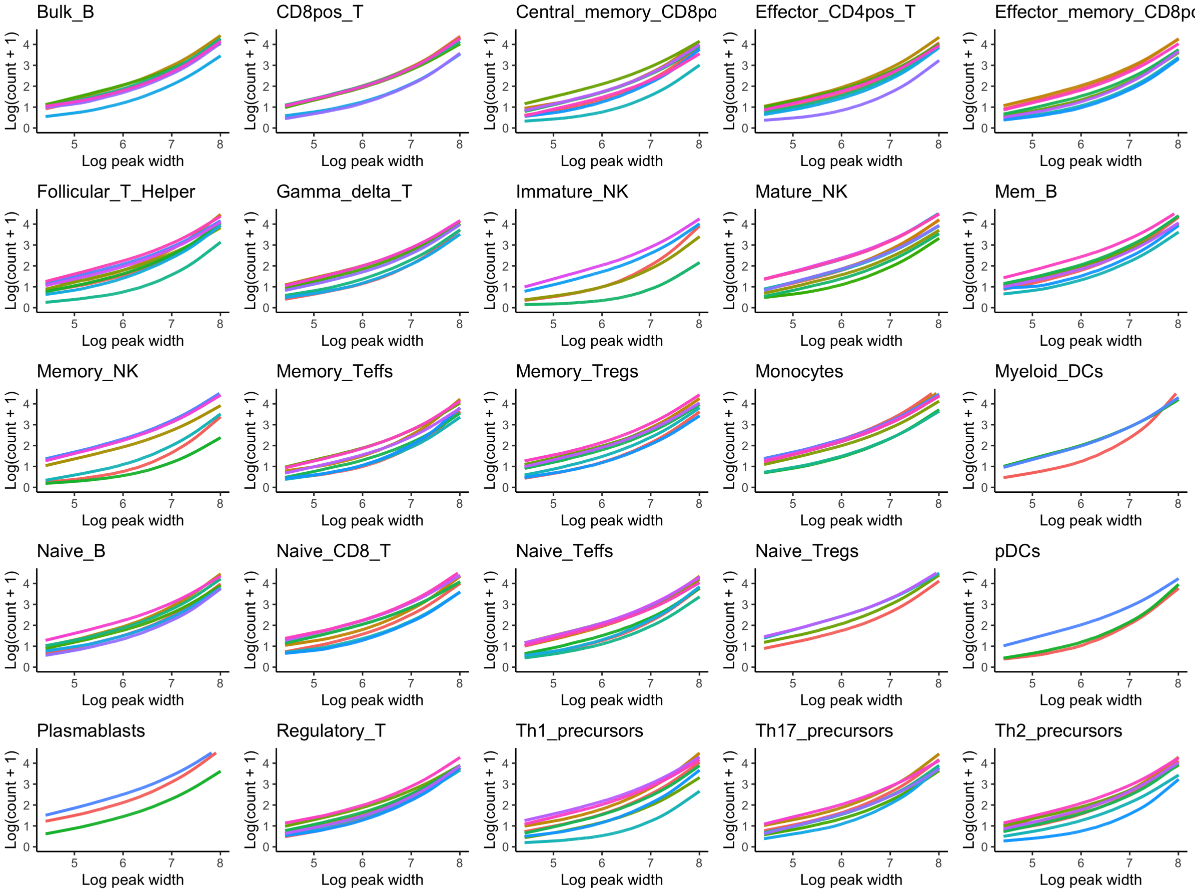


Supplementary Figure 2: *Lowess fits for the log accessibility count as a function of peak width for the dataset from Calderon et al.*[23].

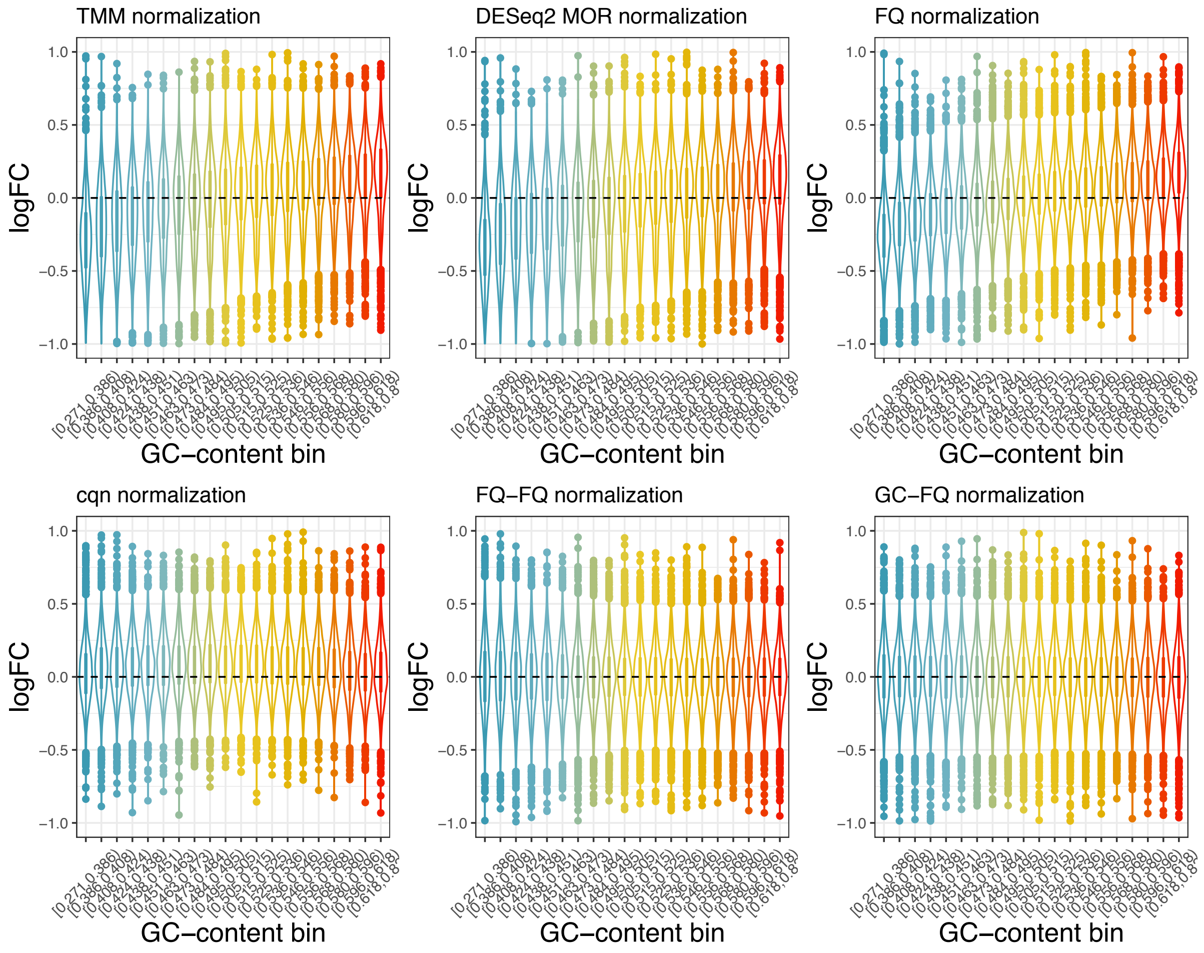


Supplementary Figure 3: *Fold-change bias for a mock comparison of stimulated mature natural killer cells in the dataset from Calderon et al.*[23].

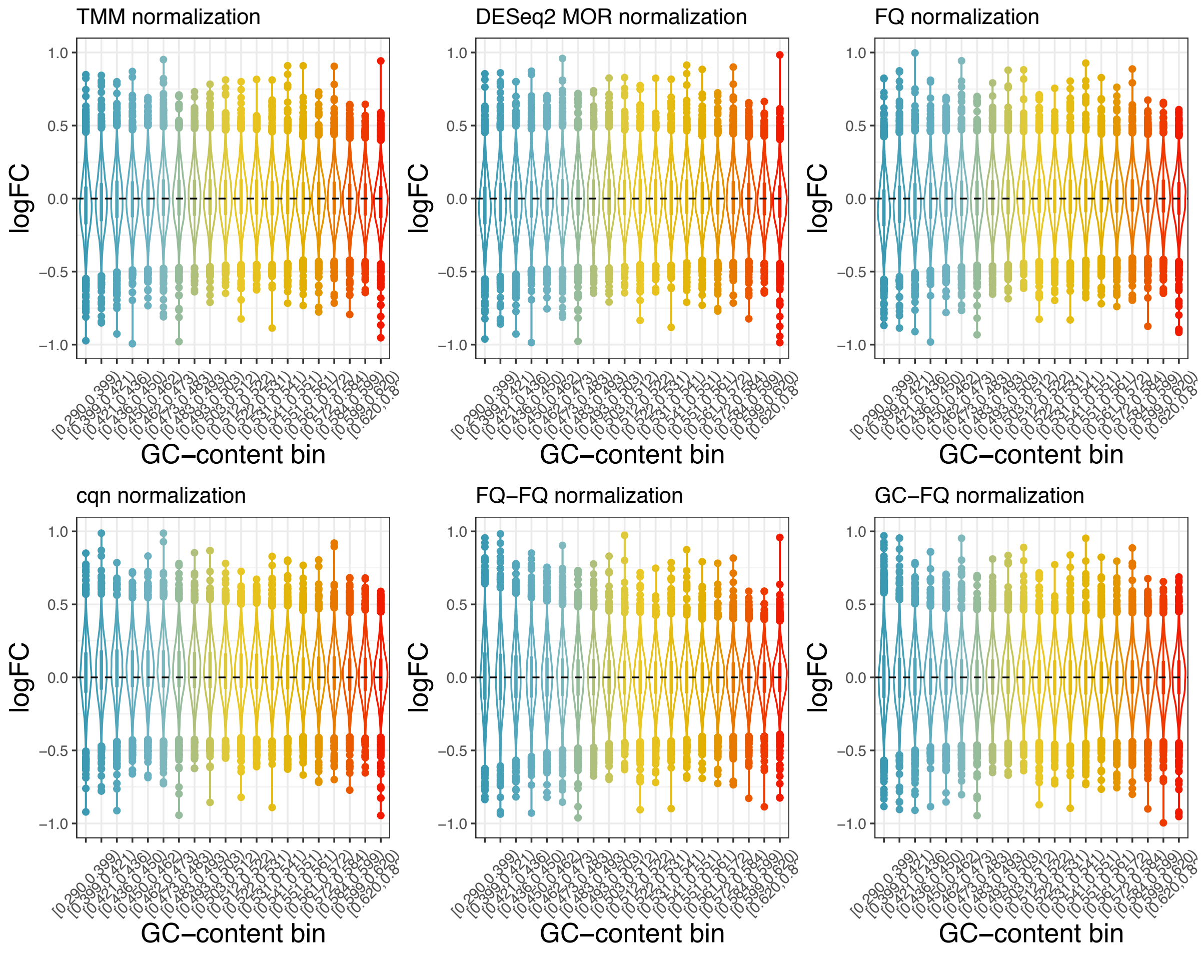


Supplementary Figure 4: *Fold-change bias for a mock comparison of stimulated nonocyte cells in the dataset from Calderon et al.*[23].

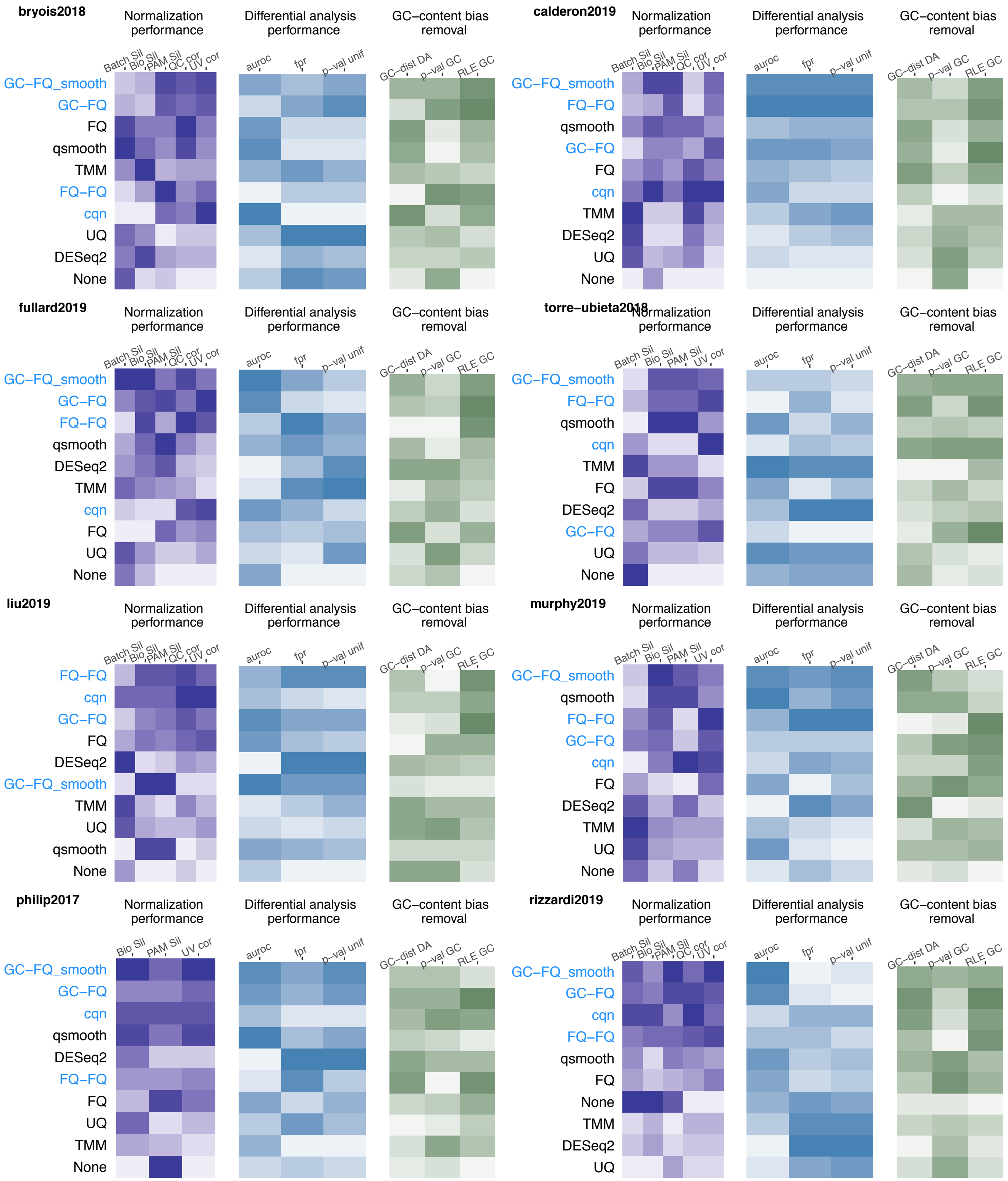


Supplementary Figure 5: *Benchmarking results for each dataset.* Each panel corresponds to the benchmarking results for each dataset, as indicated by the first author and publishing year in the top left corner. Within each panel, normalization methods are ordered from scoring well (top) to badly (bottom). The benchmark focusses on three main components: normalization performance assessment using scone, differential analysis performnance and the removal of GC-content bias, each represented by a heatmap. The pseudocolors in the heatmap represent the rank of each normalization method as compared to the other methods for that particular measure; a darker color corresponds to a better rank. All measures and normalization procedures are described in Methods. Note that not all normalization performance measures could be assessed in all datasets, since we did not have batch or QC information for some datasets.

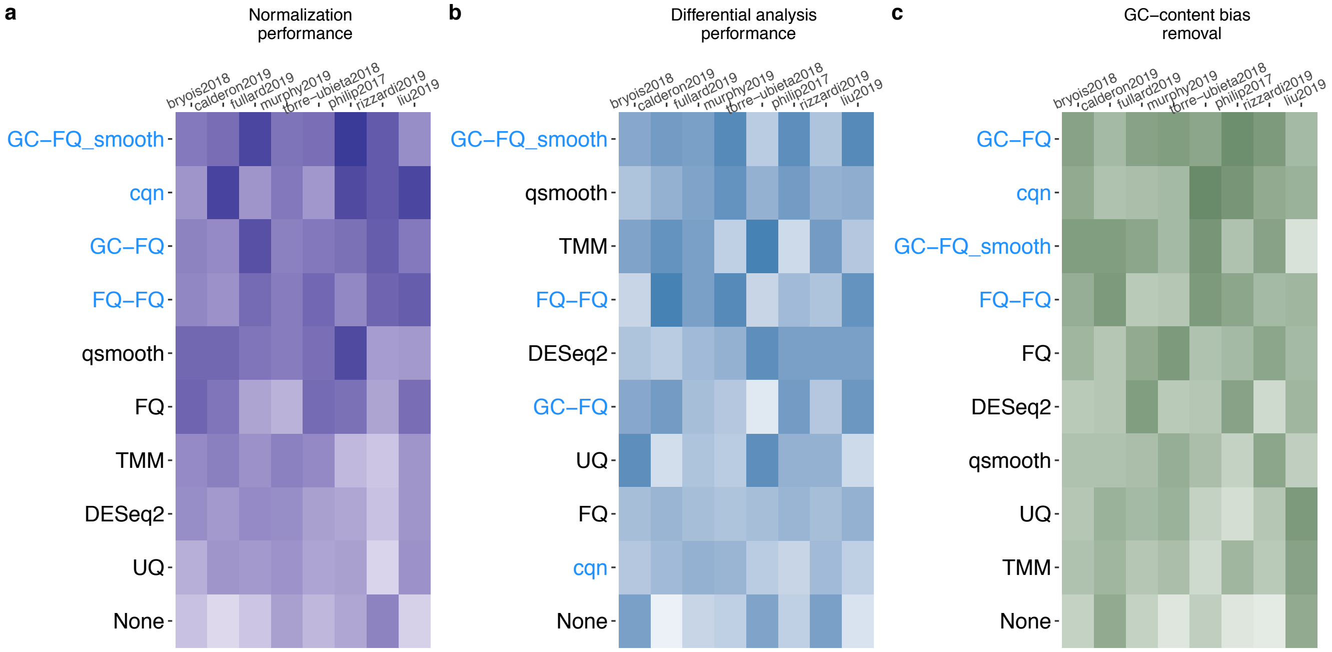


Supplementary Figure 6: *Benchmark of twelve normalization methods across eight public ATAC-seq datasets, ranked for each benchmarking component separately.* This benchmark summary shows the same results as in Figure 3, however, the methods here are ranked for each component separately. The pseudo-color images, where darker color represents better performances, display matrices of average ranks (see Methods), with rows corresponding to normalization procedures and columns to datasets. Methods are ordered according to their average rank across all evaluation criteria within each benchmarking component (i.e., separately for normalization performance, differential analysis performance and GC-content bias removal) and datasets, and their names colored based on whether they explicitly account for GC-content (blue) or not (black).

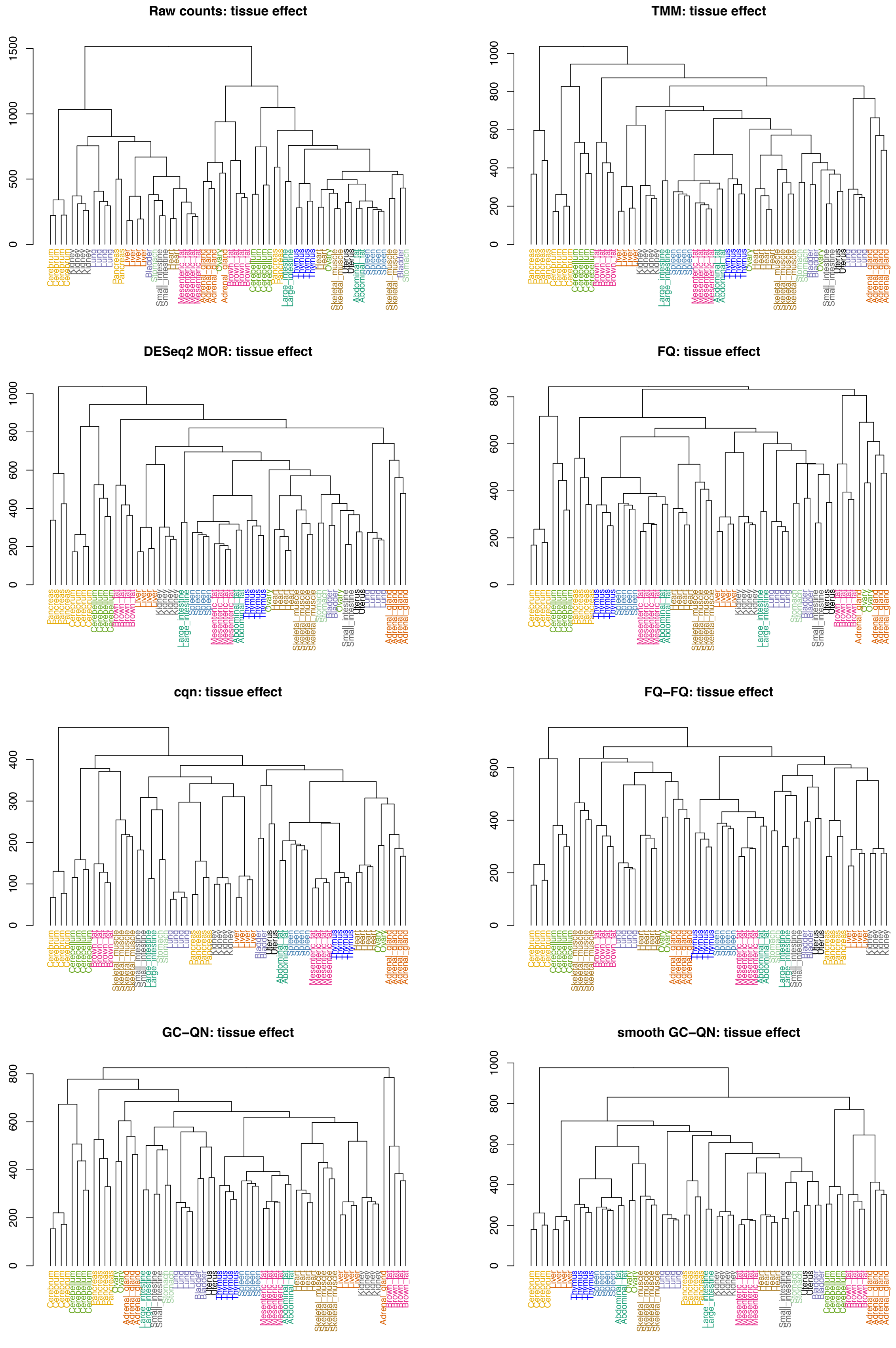


Supplementary Figure 7: *Hierarchical clustering of the dataset from Liu et al.*[30]. For each method, the normalized counts are log-transformed and pairwise Euclidean distances between samples are calculated. Agglomerative hierarchical trees are constructed using complete linkage. Samples are labeled and colored according to tissue type. The y-axis represents the Euclidean distance at which two (groups of) samples are agglomerated.

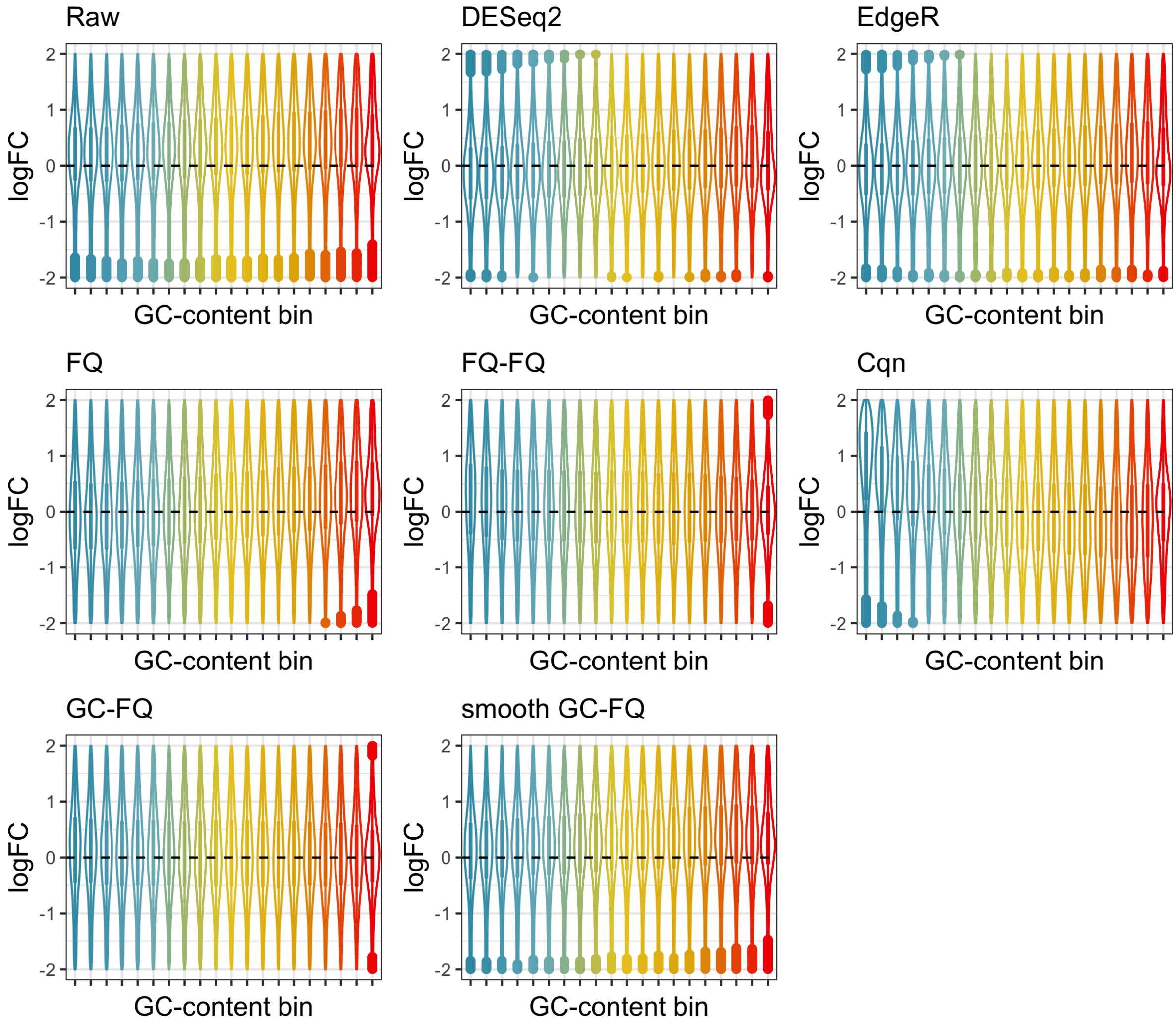


Supplementary Figure 8: *Estimated log-fold-changes for a differential accessibility analysis comparing liver with heart tissues for the dataset from Liu et al.*[30]. Each panel represents a normalization method, which was applied in its native pipeline (see Methods).

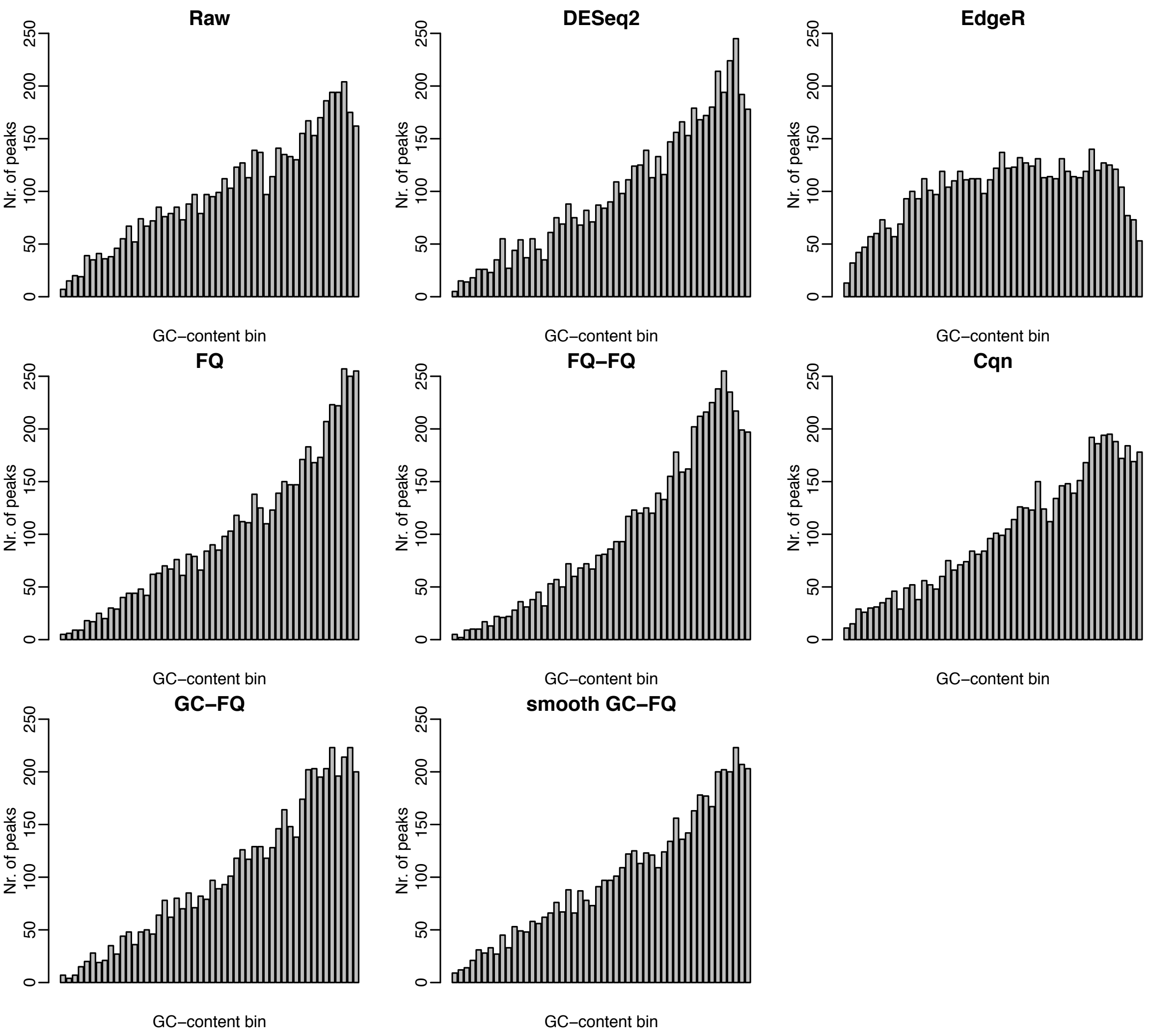


Supplementary Figure 9: *Distribution by GC-content of the top* $5,000$ *peaks for a DA analysis comparing liver with heart tissues for the dataset from Liu et al.*[30]*.* Each panel represents a normalization method, which was applied in its native pipeline (see Methods). For each normalization method, the barplots display the number of peaks (among the top $5,000$) by GC-content bin. A strong bias is observed for all methods; the most uniform distribution is seen for TMM (edgeR) normalization.

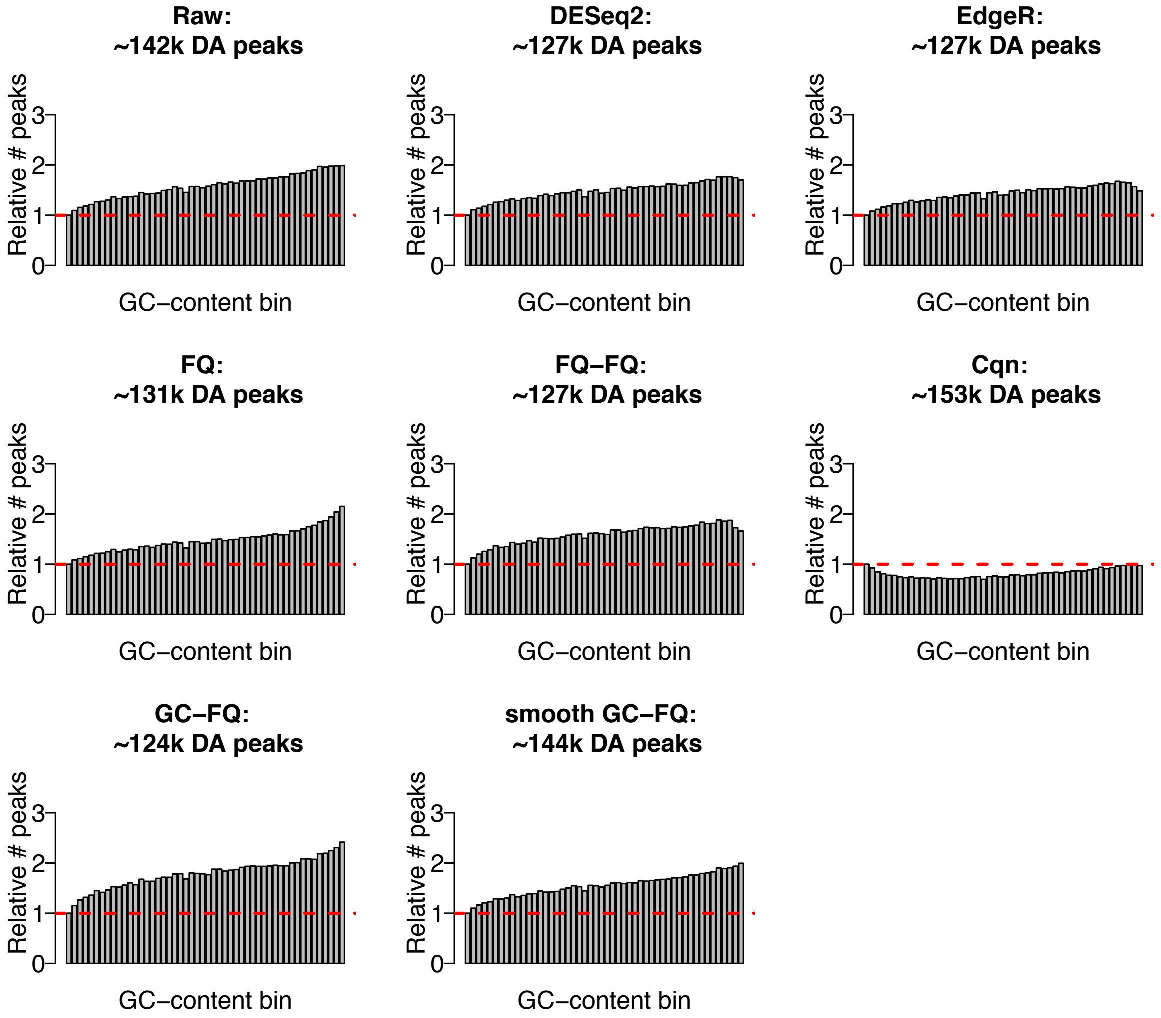


Supplementary Figure 10: *Distribution by GC-content of the significantly DA peaks for a DA analysis comparing liver with heart tissues for the dataset from Liu et al.*[30]. Each panel represents a normalization method, which was applied in its native pipeline (see Methods). For each normalization method, the barplots display the number of DA peaks at nominal FDR level of 5% relative to the corresponding number of DA peaks for the lowest GC-content bin. The total number of DA peaks found at the nominal 5% FDR level is indicated in the title of each panel.

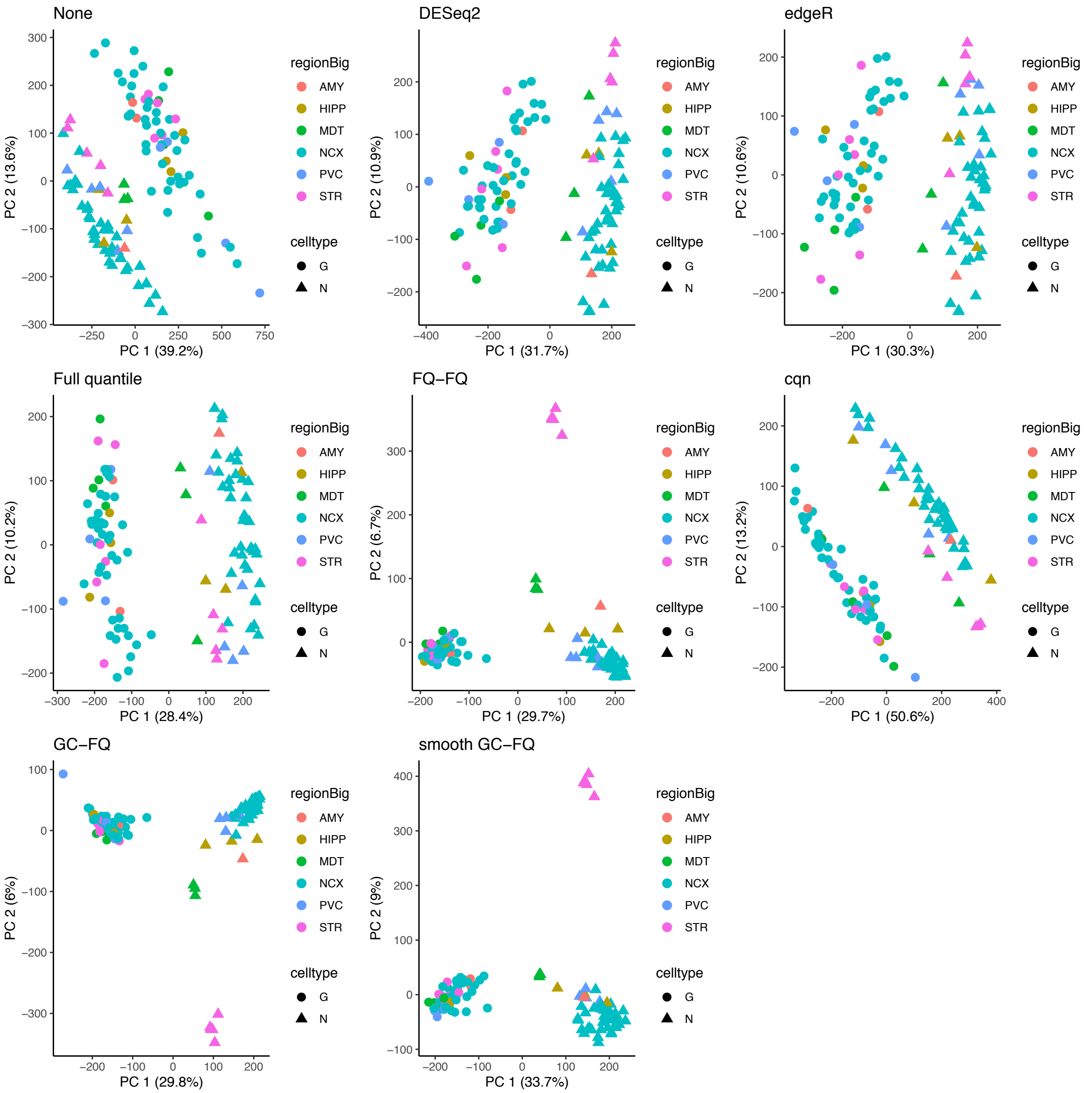


Supplementary Figure 11: *PCA plot for the dataset from Fullard et al.*[29] *after normalization.* The points are colored according to the brain region and the plotting symbol denotes the cell type.

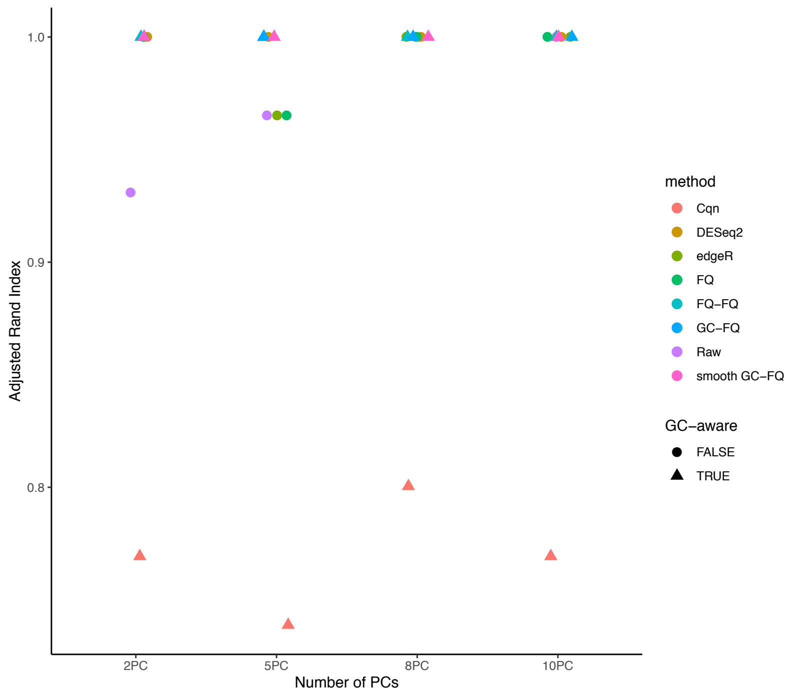


Supplementary Figure 12: *Clustering using the top principal components (PCs) for the dataset from Fullard et al.*[29]. Clustering is performed using partitioning around medoids (PAM) based on different numbers of PCs and the adjusted Rand index (ARI) is calculated with respect to the true cell type labels. All methods except `cqn` correctly recover the two cell types, resulting in an ARI value of 1.

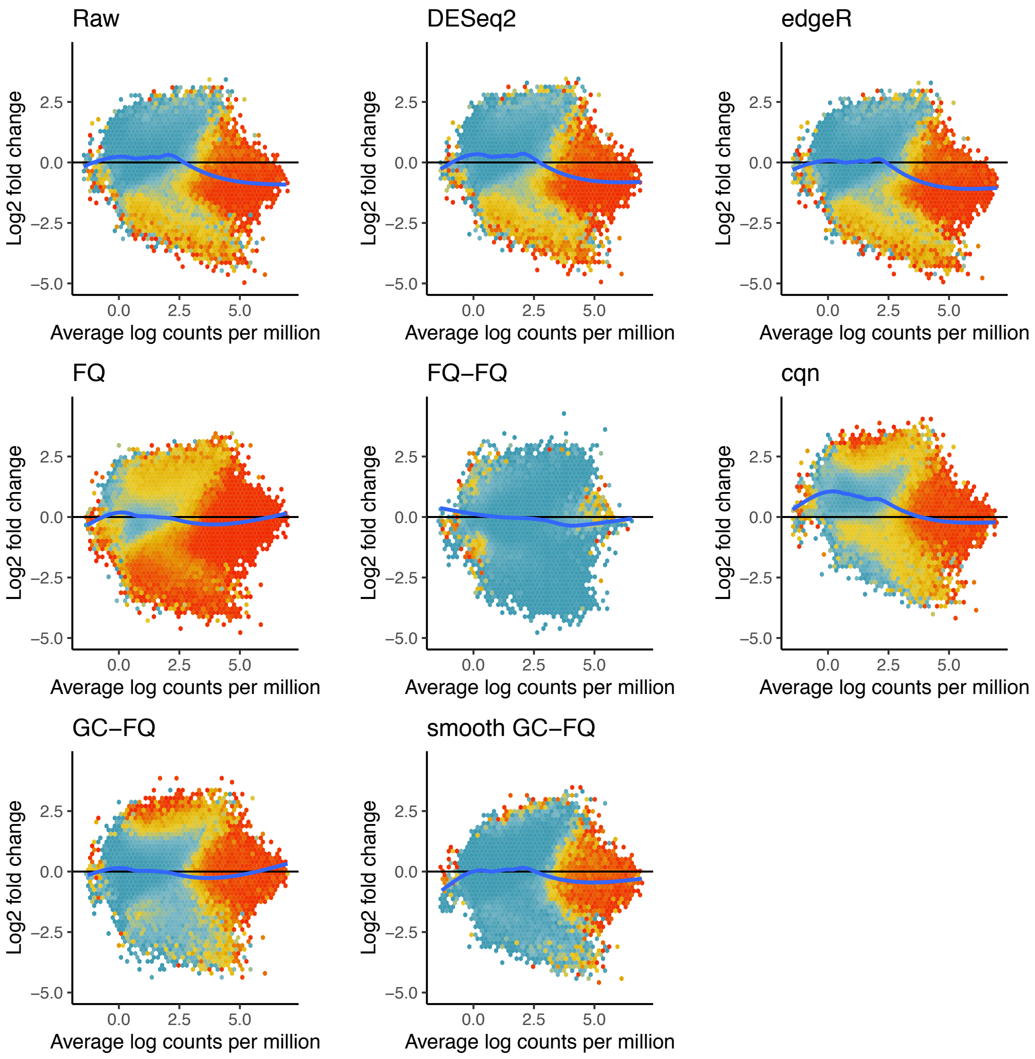


Supplementary Figure 13: *MD-plots with hexagonal binning comparing neuronal to non-neuronal cells for the dataset from Fullard et al.*[29]. The x-axis represents average log-count per million and the y-axis represents log-fold-change as estimated based on a negative binomial model fitted using edgeR (or DESeq2 for DESeq2 normalization). Each panel corresponds to a normalization method. The hexagons are colored according to the average GC-content of the corresponding peaks, where red denotes high GC-content and blue denotes low GC-content.

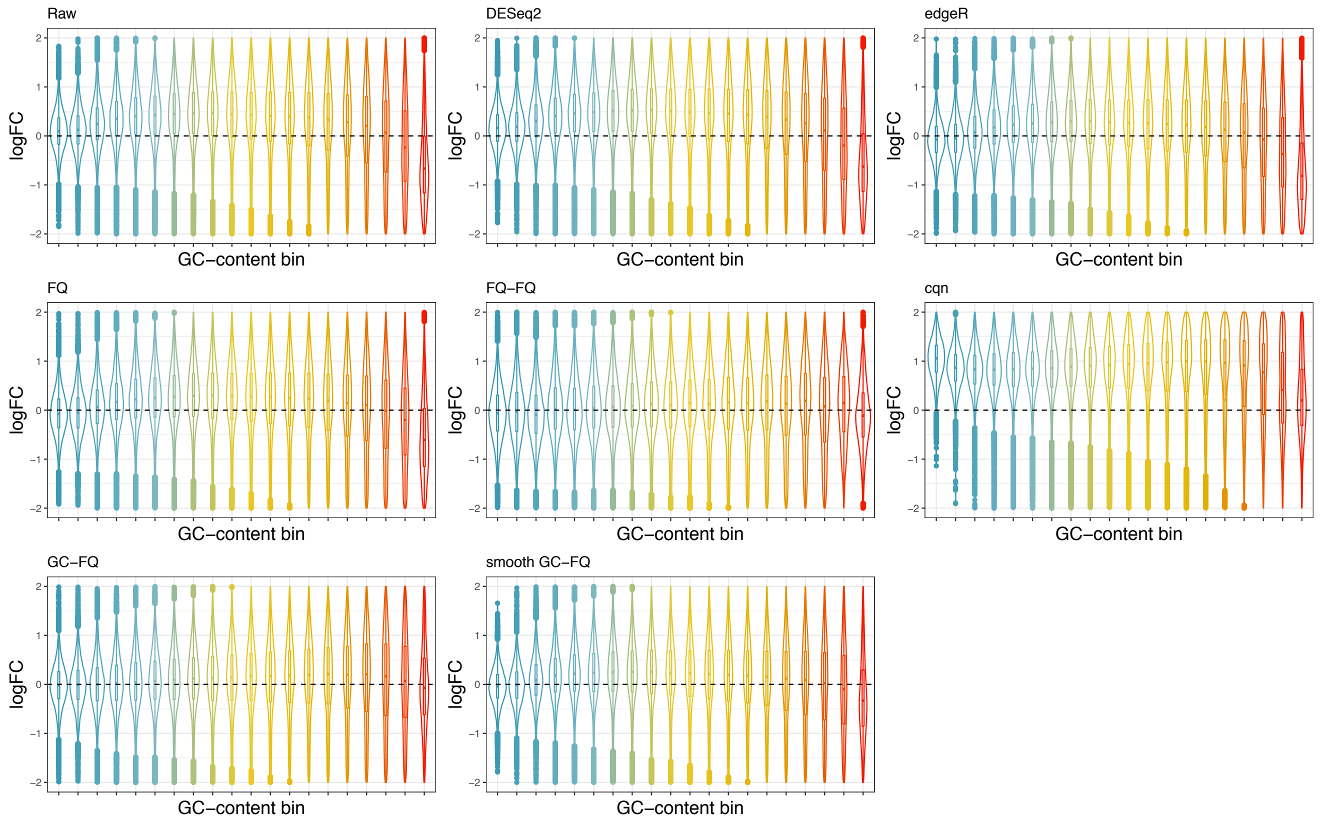


Supplementary Figure 14: *Stratified violin plots of log-fold-changes by GC-content for the dataset from Fullard et al.* [29]. The x-axis represents equally-sized GC-content bins and the y-axis represents log-fold-change as estimated based on a negative binomial model fitted using edgeR. Each panel corresponds to a normalization method.

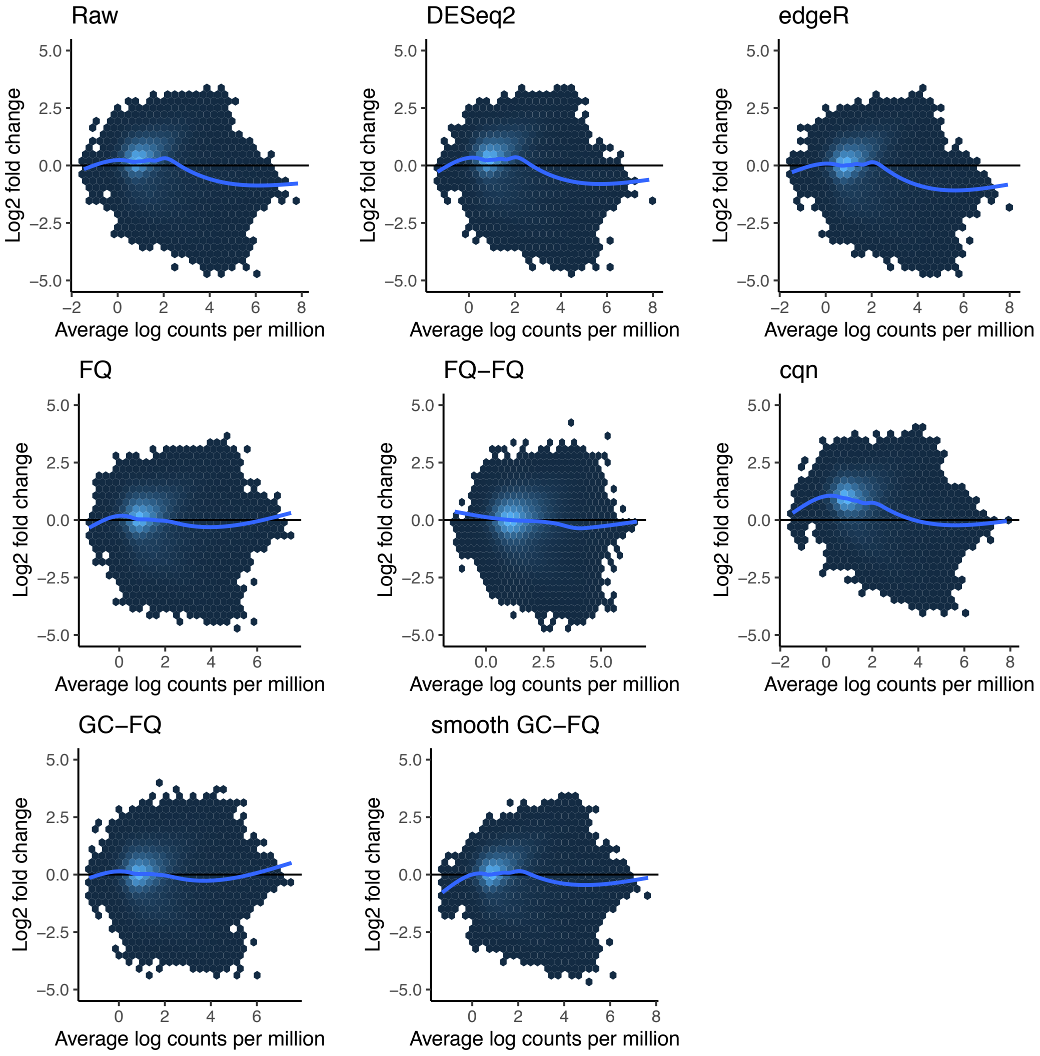


Supplementary Figure 15: *MD-plots with hexagonal binning comparing neuronal to non-neuronal cells for the dataset from Fullard et al.*[29]. The x-axis represents average log-count per million and the y-axis represents log-fold-change as estimated based on a negative binomial model fitted using `edgeR` (or `DESeq2` for `DESeq2` normalization). Each panel corresponds to a normalization method. The hexagons are colored according to the number of peaks in each hexagon, with lighter colors corresponding to more peaks, and darker colors to less peaks.

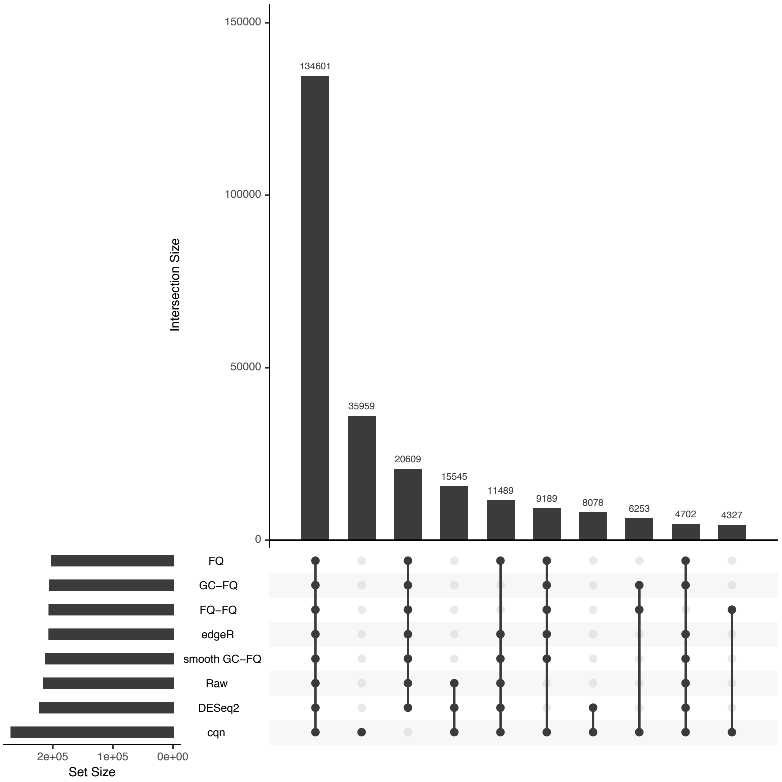


Supplementary Figure 16: *UpSet plot comparing overlap in DA peaks for different normalization procedures for the dataset from Fullard et al.*[29]*.* Left panel: Barplot of the number of DA peaks for each normalization method (5% nominal FDR level). Top panel: Barplot of the number of DA peaks in common for every combination of methods. Center panel: Combination of methods under consideration for the top panel. For example, the first column shows that $134,601$ DA peaks are found by all methods.

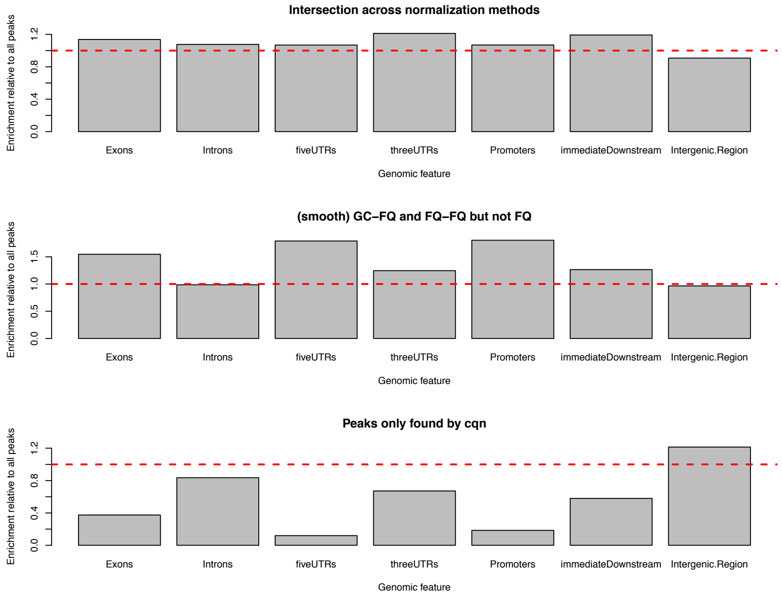


Supplementary Figure 17: *Enrichment of genomic features relative to background for the dataset from Fullard et al.* [29]. Upper panel: Barplot of the enrichment of each genomic feature relative to all peaks in the dataset, for the intersection of peaks discovered by all normalization methods. An enrichment of relevant genomic features, such as promoters, 5′ UTRs, and exons is observed. Middle panel: Similar, for peaks discovered by FQ-FQ and (smooth) GC-FQ but not FQ, where substantial enrichment of promoters, 5′ UTRs, and exons is observed. Bottom panel: Similar, for peaks only discovered by cqn normalization. No enrichment of relevant features is observed.

### 4 Supplementary Tables

Supplementary Table 1: *Gene Ontology analysis of DA peaks.* Top 20 enriched Biological Process (BP) Gene Ontology terms for the DA peaks identified by all normalization procedures, when comparing neuronal vs. non-neuronal cells for the BOCA dataset from Fullard et al.[29]. A total of 68 GO terms were significantly enriched at a 5% nominal FDR level.

| | GO ID | GO term | BH-adjusted *p*-value |
|---|---|---|---|
| 1 | GO:0007399 | nervous system development | 0.0000 |
| 2 | GO:0022008 | neurogenesis | 0.0000 |
| 3 | GO:0048666 | neuron development | 0.0001 |
| 4 | GO:0030182 | neuron differentiation | 0.0001 |
| 5 | GO:0031175 | neuron projection development | 0.0001 |
| 6 | GO:0048699 | generation of neurons | 0.0001 |
| 7 | GO:0045664 | regulation of neuron differentiation | 0.0005 |
| 8 | GO:0032990 | cell part morphogenesis | 0.0005 |
| 9 | GO:0050767 | regulation of neurogenesis | 0.0005 |
| 10 | GO:0007417 | central nervous system development | 0.0005 |
| 11 | GO:0031329 | regulation of cellular catabolic process | 0.0007 |
| 12 | GO:0051960 | regulation of nervous system development | 0.0007 |
| 13 | GO:0030030 | cell projection organization | 0.0007 |
| 14 | GO:0000122 | negative regulation of transcription by RNA polymerase II | 0.0007 |
| 15 | GO:0007420 | brain development | 0.0007 |
| 16 | GO:0030900 | forebrain development | 0.0007 |
| 17 | GO:0120036 | plasma membrane bounded cell projection organization | 0.0007 |
| 18 | GO:0098732 | macromolecule deacylation | 0.0008 |
| 19 | GO:0048812 | neuron projection morphogenesis | 0.0008 |
| 20 | GO:0009894 | regulation of catabolic process | 0.0008 |

Supplementary Table 2: *Gene Ontology analysis of DA peaks.* Top 20 enriched Biological Process Gene Ontology terms for DA peaks discovered by FQ-FQ and (smooth) GC-FQ but not by FQ normalization, when comparing neuronal vs. non-neuronal cells for the BOCA dataset from Fullard et al.[29]. No GO terms were significantly enriched at a 5% nominal FDR level.

| | GO ID | GO term | BH-adjusted $p$-value |
|---|---|---|---|
| 1 | GO:0050803 | regulation of synapse structure or activity | 0.0965 |
| 2 | GO:0032922 | circadian regulation of gene expression | 0.0965 |
| 3 | GO:0007416 | synapse assembly | 0.0965 |
| 4 | GO:0050808 | synapse organization | 0.0965 |
| 5 | GO:0001964 | startle response | 0.1206 |
| 6 | GO:0050807 | regulation of synapse organization | 0.1484 |
| 7 | GO:0060291 | long-term synaptic potentiation | 0.2113 |
| 8 | GO:0099536 | synaptic signaling | 0.2113 |
| 9 | GO:0007268 | chemical synaptic transmission | 0.2113 |
| 10 | GO:0098916 | anterograde trans-synaptic signaling | 0.2113 |
| 11 | GO:0097120 | receptor localization to synapse | 0.2113 |
| 12 | GO:0051963 | regulation of synapse assembly | 0.2113 |
| 13 | GO:0007623 | circadian rhythm | 0.2113 |
| 14 | GO:0051965 | positive regulation of synapse assembly | 0.2113 |
| 15 | GO:0099537 | trans-synaptic signaling | 0.2113 |
| 16 | GO:0099632 | protein transport within plasma membrane | 0.2113 |
| 17 | GO:0099637 | neurotransmitter receptor transport | 0.2113 |
| 18 | GO:0050806 | positive regulation of synaptic transmission | 0.2372 |
| 19 | GO:0050804 | modulation of chemical synaptic transmission | 0.2372 |
| 20 | GO:0033555 | multicellular organismal response to stress | 0.2372 |

Supplementary Table 3: *Gene Ontology analysis of DA peaks.* Top 20 enriched Biological Process (BP) Gene Ontology terms for the DA peaks uniquely identified by *cqn*, when comparing neuronal vs. non-neuronal cells for the BOCA dataset from Fullard et al.[29]. No GO terms were significantly enriched at a 5% nominal FDR level.

| | GO ID | GO term | BH-adjusted $p$-value |
|---|---|---|---|
| 1 | GO:0010842 | retina layer formation | 0.1253 |
| 2 | GO:0072080 | nephron tubule development | 0.1253 |
| 3 | GO:0060993 | kidney morphogenesis | 0.1253 |
| 4 | GO:0061326 | renal tubule development | 0.1253 |
| 5 | GO:0072078 | nephron tubule morphogenesis | 0.1314 |
| 6 | GO:0072088 | nephron epithelium morphogenesis | 0.1314 |
| 7 | GO:0061333 | renal tubule morphogenesis | 0.1314 |
| 8 | GO:0072028 | nephron morphogenesis | 0.1314 |
| 9 | GO:0072009 | nephron epithelium development | 0.1314 |
| 10 | GO:0060675 | ureteric bud morphogenesis | 0.2517 |
| 11 | GO:0001657 | ureteric bud development | 0.2517 |
| 12 | GO:0010874 | regulation of cholesterol efflux | 0.2517 |
| 13 | GO:0072171 | mesonephric tubule morphogenesis | 0.2517 |
| 14 | GO:0072163 | mesonephric epithelium development | 0.2517 |
| 15 | GO:0072164 | mesonephric tubule development | 0.2517 |
| 16 | GO:0010985 | negative regulation of lipoprotein particle clearance | 0.2711 |
| 17 | GO:0001823 | mesonephros development | 0.2796 |
| 18 | GO:0072073 | kidney epithelium development | 0.2881 |
| 19 | GO:0072006 | nephron development | 0.2987 |
| 20 | GO:0048593 | camera-type eye morphogenesis | 0.3765 |
